# Polar accumulation of pyoverdin and exit from stationary phase

**DOI:** 10.1101/2021.07.27.453990

**Authors:** Clara Moreno-Fenoll, Maxime Ardré, Paul B. Rainey

## Abstract

Pyoverdin is a water-soluble metal-chelator synthesized by members of the genus *Pseudomonas* and used for the acquisition of insoluble ferric iron. Although freely diffusible in aqueous environments, preferential dissemination of pyoverdin among adjacent cells, fine-tuning of intracellular siderophore concentrations, and fitness advantages to pyoverdin-producing versus nonproducing cells, indicate control of location and release. Here, using time-lapse fluorescence microscopy to track single cells in growing microcolonies of *Pseudomonas fluorescens* SBW25, we show accumulation of pyoverdin at cell poles. Accumulation occurs on cessation of cell growth, is achieved by cross-feeding in pyoverdin-nonproducing mutants and is reversible. Moreover, accumulation coincides with localization of a fluorescent periplasmic reporter, suggesting that pyoverdin accumulation at cell poles is part of the general cellular response to starvation. Compatible with this conclusion is absence of non-accumulating phenotypes in a range of pyoverdin mutants. Analysis of the performance of pyoverdin-producing and nonproducing cells under conditions promoting polar accumulation shows an advantage to accumulation on resumption of growth after stress. Examination of pyoverdin polar accumulation in a multispecies community and in a range of laboratory and natural species of *Pseudomonas*, including *P. aeruginosa* PAO1 and *P. putida* KT2440, confirms that the phenotype is characteristic of *Pseudomonas*.

## Introduction

Extracellular secreted products perform important functions in microbial populations and communities. They provide structure and protection [1], enable coordinated action [2] [3] and also allow acquisition of recalcitrant nutrients, such as polymers that are too large to be internalized [4], or are otherwise unavailable. An example of the latter is pyoverdin.

Pyoverdin is a naturally fluorescent iron-scavenging chelator (sideropohore) produced by members of the genus *Pseudomonas* [5]. Iron is an essential micronutrient that exists in an insoluble state (ferric) in aerobic environments [6]. In response to intracellular iron scarcity, pyoverdin biosynthesis begins in the cytoplasm via nonribosomal peptide synthesis and undergoes maturation in the periplasm [7]. After secretion, it binds Fe^3+^ with high affinity (Ka = 10^32^ M^-1^ for PVDI produced by *P. aeruginosa* PAO1 [8]). Once bound to ferric iron the ferripyoverdin complex loses its fluorescent properties, but is recognized by a specific receptor (FpvA) and imported back into the periplasm, where iron is extracted. The pyoverdin molecule is then recycled and can undergo further cycles of export and import [9].

Because pyoverdin is a soluble extracellular product, it has been widely assumed to be equally available to all members of a community [10, 11, 12]. However, recent work shows that its distribution is subject to cell-level control. In one study pyoverdin producers retained an environment-dependent fitness advantage in conditions where invasion by nonproducers was expected. In light of these experimental results, the possibility of personalization was raised [13]. Other studies have further supported this notion. For example, in growing microcolonies, pyoverdin diffuses primarily through adjacent cells, reducing loss into the environment [14]. Furthermore, *Pseudomonas aeruginosa* cells tune periplasmic concentrations of pyoverdin in order to protect against oxidative stress [15].

Here we use time-lapse fluorescence microscopy to study the relationship between *P. fluorescens* SBW25 cells and pyoverdin. Recognizing the importance of spatial structure and contributions therefrom to *µ*m-scale features of microbial assemblages [16, 17, 18, 19], cells were grown on thin layers of agarose set on top of microscope slides. In actively dividing cells, naturally fluorescent apo-pyoverdin is evenly distributed in the periplasm, however, on cessation of growth we observed pyoverdin to accumulate at cell poles. This surprising discovery motivated quantitative analysis, revealing the process of accumulation to be dynamic, reversible, and ecologically relevant.

## Results

*P. fluorescens* SBW25 (hereafter SBW25) is a model bacterium [20] known to produce pyoverdin [21] and other secreted products [22, 23]. In a previous study Zhang & Rainey [13] provided evidence of pyoverdin personalization that was inferred following contrasting outcomes of fitness assays performed under different conditions. We reproduced these assays, but rather than examining frequencies of producers and nonproducers by plating, aliquots were observed by fluorescence microscopy. In casamino acids medium (CAA), the medium where an unexpected advantage for pyoverdin producers had been described, cells exhibited internal pyoverdin accumulation more evident at the pole where fluorescent foci (Fig. 1 A) appeared. This initial observation provoked further analysis.

**Fig. 1.**
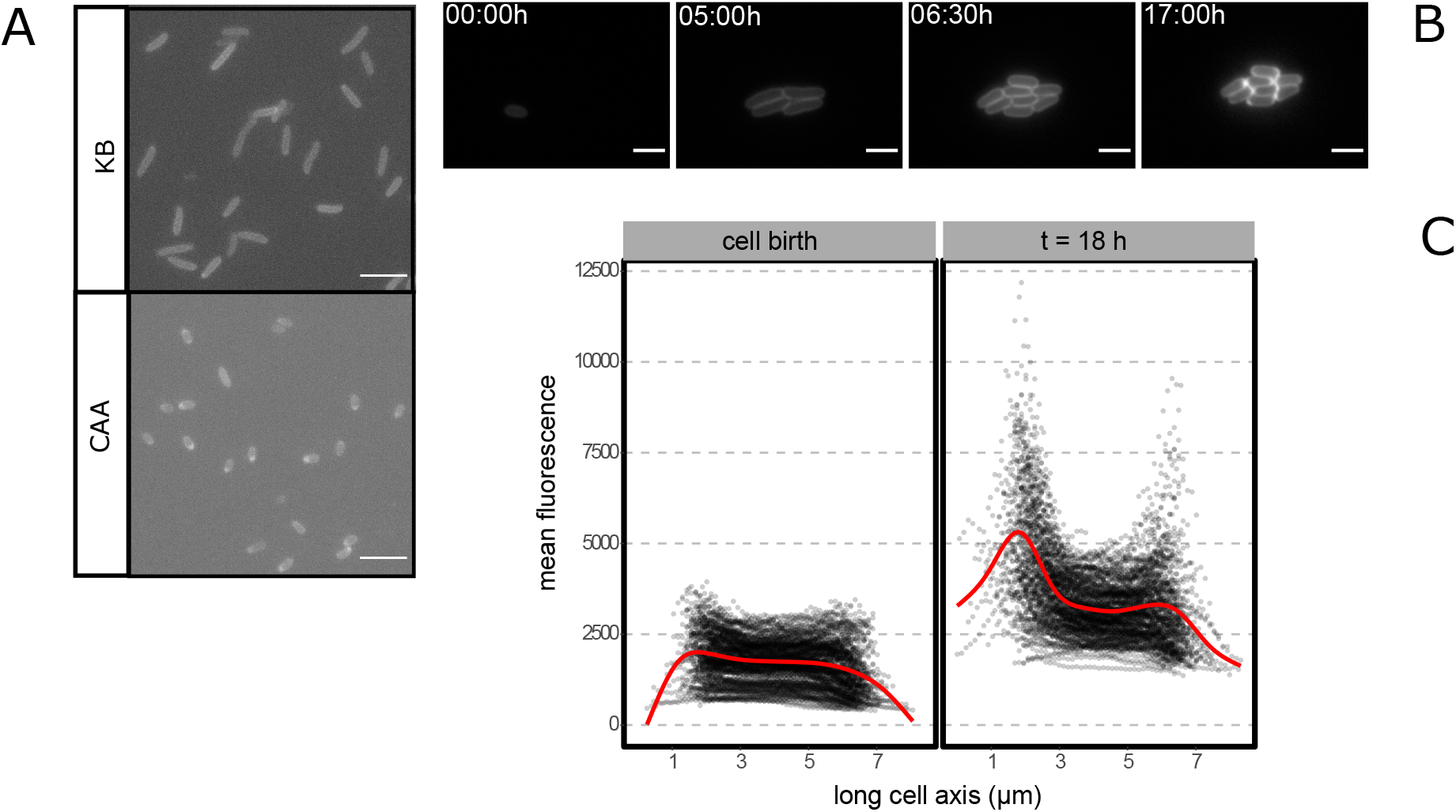
Accumulation of pyoverdin in *P. fluorescens* SBW25 at the cell pole. A) Snapshots of experiments described in [13] where fitness assays of pyoverdin producing SBW25 and a nonproducing *pvdS* defective mutant yield contrasting results depending on the culture medium. In both cases the environment is unstructured and ancestral SBW25 producer cells are rare, inoculated at 1%. A fitness advantage to nonproducing cells in KB was previously reported, but the reverse in CAA [13]. In KB (top) pyoverdin nonproducing cells rarely showed evidence of accumulation of pyoverdin, whereas (bottom) this was common in CAA cultured cells where accumulation is visible at the pole. Images were obtained from 3 *µ*l samples of these experiments imaged under fluorescence light to visualize the distribution of pyoverdin. All scale bars correspond to 10 *µ*m. B) Fluorescence time-lapse images of a growing microcolony of SBW25 in a SMM agarose pad. Images represent selected time points including, respectively: the initial inoculum, exponential growth, end of exponential growth (i.e., the final number of cells in the colony) and end of time-lapse acquisition (18 h total) C) Mean fluorescence intensity along the long axis of cells in a growing microcolony, when the last generation of cells is born (left) and at the end of acquisition (t = 18 h, right). Black dotted line represents the fluorescence profile of individual cells, the red line represents a smoothed mean of all the cells. Between 0 h and 18 h accumulation of pyoverdin is evident, especially at the pole.

To accurately characterize subcellular patterns of pyoverdin, time-lapse fluorescence images of ancestral SBW25 were obtained in defined succinate minimal medium (SMM), where succinate acts both as a carbon source and a weak iron chelator. During exponential phase and early stationary phase, SBW25 cells showed a phenotype typical of fluorescent *Pseudomonas* with pyoverdin being homogeneously distributed in the periplasm, while in late stationary phase fluorescent pyoverdin foci appeared at the cell pole (Fig. 1 B). Superposing cell fluorescence profiles obtained by image segmentation confirmed that both polar localization and accumulation of the siderophore occurred at this later stage (Fig. 1 C).

By examining cell division throughout the time-lapse series it is possible to explore the relationship between time and population growth. The corresponding frequency of polarized cells (Fig. 2 A) was tracked by classifying segmented cells automatically as “polarized” (accumulated) or “homogeneous” (non-accumulated) using a machine learning algorithm (Supplementary Materials and Methods). After inoculating the microscope slide, cells undergo a period of acclimation to the medium without division. We observed no accumulation events during this phase. As division begins, age of cells in microcolonies decreases until it reaches a minimum that marks exponential phase. Pyoverdin continued to be non-polarized. Note that the small frequency of cells identified as “polarized” in the plot falls within the range of classification error. Finally, the population enters stationary phase and cells continue ageing without division. Accumulation of pyoverdin at the pole increased within the first few hours, encompassing most cells within the population by 18h (Fig. S1). As stationary phase progresses, cells become classified as “polarized” at different rates while possessing varying amounts of intracellular pyoverdin, suggesting that the quantity of internal pyoverdin does not define polar localization per se (Fig S2 A). No significant difference was observed in the total amount of internal pyoverdin between cells classified as “polarized” or “homogeneous” (Fig S2B), despite the fact that the majority of cells accumulate pyoverdin over time (Fig 1C).

**Fig. 2.**
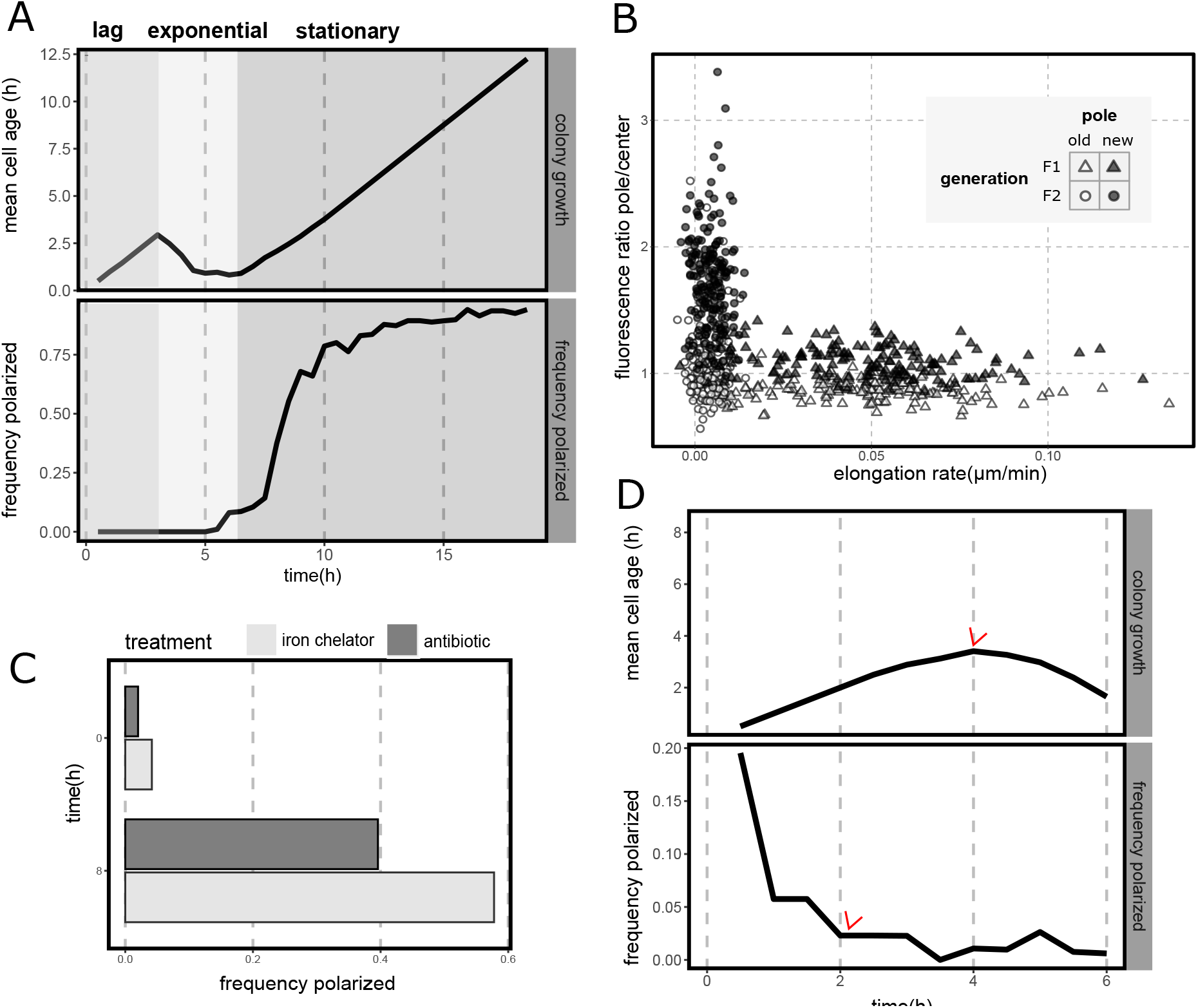
Polar accumulation is a reversible phenotype associated with arrest of cell division. A) Polarization in different growth stages of a microcolony. Mean age of the cells in a growing microcolony of SBW25 (black line, top) and the corresponding frequency of polarized cells for each time point (black line, bottom). Colored panels represent the growth stages of the microcolony, from left to right: lag phase (generation F0), exponential phase (generation F1), and stationary phase (generation F2). N = 35, 159, 187 respectively. In all cases data has been filtered to exclude cells with segmentation errors or other artefacts that preclude proper analysis. Note that as microcolonies begin to form in exponential phase and cells are no longer isolated, overlap between adjacent cells creates regions of high fluorescence that could lead to classification errors. Nevertheless, visual inspection reveals that cells remain in a homogeneous state during exponential growth, with polarization onset being clearly associated to stationary phase. To maintain a consistent physiological response, in this study cells are manipulated from a starting point corresponding to early stationary phase (Fig. S4. B) Elongation rate and accumulation of fluorescence at the cell pole of individual cells in a growing microcolony. Data extracted from A). Markers represent individual cells in different growth phases of the colony (F1, exponential, triangle markers; F2, stationary, circle markers) and the old (white markers) and new (black markers) cell pole. Elongation rate is represented by the average over the lifetime of a cell. Accumulation of fluorescence at the pole is represented by the maximum ratio over the lifetime of a cell of the sum of the pixels in the pole region and central region of a cell. These regions are defined by segmenting the cell and dividing it in 3 portions over the long axis, where the external 1/3 represent each pole and the remaining 1/3 represents the center. C) Polarization in response to chemical stresses related and unrelated to iron metabolism. Bars represent the frequency of polarized cells at the start of treatment and after 8h of treatment with either 100 *µ*g/ml 2,2’-dipyridil (DP) (light bars) or 5 *µ*g/ml tetracycline (dark bars). No cell division was observed during treatment. N = 99 (t=0h), 94 (t = 8h) and N = 94 (t = 0h), 79 (t = 8h) for DP and tetracyline treatments respectively. D) Depolarization and subsequent growth of cells pre-treated with an iron chelator. Plot represents colony growth and polarization as in A). Cells were treated with 100 *µ*g/ml DP during 4h, washed and inoculated on a fresh SMM agarose pad. Data corresponds to five technical replicates i.e., five positions on the agarose pad. Red arrows indicate the time point where the colony overall starts growing (top) and where the majority of the cells are depolarized (bottom). Total initial number of cells N=87.

While the time-averaged state of microcolonies exposes population-level dynamics of polarization – namely, that accumulation happens in stationary phase – it obscures the behavior of individual cells. To specifically analyze single cells, cells were grouped according to division status over three generations: F0 for initial inoculum (cells where birth could not be identified but division was observed), F1 for cells in exponential phase (born from the division of F0 cells and underwent division later) and F2 for the daughter cells of F1 that entered stationary phase and remained constant for the final hours of the experiment. Growth measurements from F1 cells, and pyoverdin accumulation measurements from F2 cells, corroborate – despite some variability in the onset of accumulation – that accumulation is incompatible with active cell division. Cells either elongate or accumulate fluorescence (pyoverdin) at the cell pole. This result holds for both the old and new pole (Fig. 2 B). Curiously, pyoverdin accumulates preferentially at the new pole, but not exclusively, and sometimes distinct foci are present at both (Fig. S3)

Stationary phase marks cessation of cell division due to nutrient depletion. On an agarose pad, entry into stationary phase is highly variable, with access of cells to nutrients and oxygen depending on position and proximity to neighbouring cells. To study pyoverdin accumulation under controlled conditions cells were exposed to two stressors that abruptly arrest cell division, but by different mechanisms: 1, an iron chelating agent (2,2’-dipyridil (DP)) and; 2, a protein synthesis-disrupting antibiotic (tetracycline)(Fig. S5 A). Both stressors induced accumulation at the pole (Fig. 2 C).

Precisely because polarization appears when cells are starved and/or stressed, processes associated with cell death, such as cell wall damage, are potential elicitors. We thus asked whether pyoverdin accumulation was connected to cell viability. Bacteria were treated with DP, which reliably induces polarization (Fig. 2 C) and then transferred to a fresh agarose pad. Time-lapse imaging revealed that polarization is reversible and that depolarization precedes exit from lag phase (Fig. 2 D). Polarized cells consistently re-established a homogeneous pyoverdin distribution within the first few hours after inoculation, with elongation and division resuming at a later time point (Fig. S5 B). Note that both the initial frequency of polarized cells and the time to depolarization were counted from the start of image acquisition. During the previous experimental manipulation, some individuals might have altered their polarization status. Despite this, the trend is clear, with depolarization occurring before resumption of growth. Polarization is thus not a consequence of cell death but rather a reversible, dynamic process tied to arrest of cell division.

Pyoverdin polarization is clearly connected to the physiological state of cells, but how this is achieved is unclear. A recent study showed that the cytoplasm of *E*.*coli* shrinks on starvation leading to enlargement of periplasmic space at cell poles [24]. We hypothesized that this might also occur in SBW25 cells, with the space created by shrinkage allowing accumulation of pyoverdin. To this end, a red fluorescent protein (mScarlet) was translationally fused to the peptide signal from *dsbA* thus targeting mScarlet to the periplasm [24]. Having confirmed periplasmic localization, cells were grown on agarose pads and the distribution of mScarlet was tracked by microscopy. At inoculation, mScarlet was homogeneously distributed as observed for pyoverdin, however, on entry into stationary phase, the fluorescent marker accumulated at the poles of the cells (Fig. 3 A). As with pyoverdin, mScarlet accumulation was biased toward new poles (Fig. 3 B). As expected, unlike pyoverdin, mScarlet showed no evidence of secretion to the external medium (Fig. S8)

**Fig. 3.**
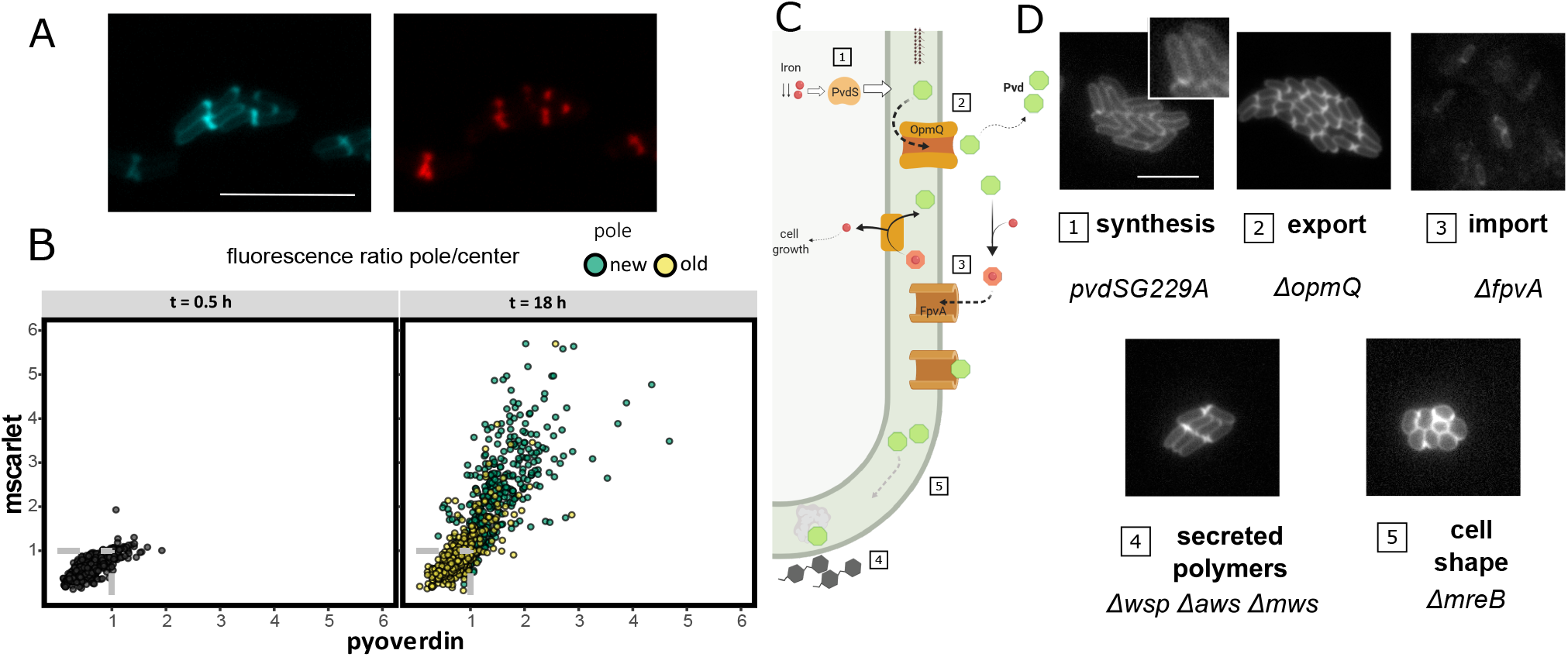
Untangling the mechanism of polarization. A) A recent study [24] showed that upon starvation the cytoplasm of *E. coli* cells shrinks creating extra space in the periplasm at one of the cell poles. To check if a similar phenomenon happens in *P. fluorescens* SBW25 we expressed a red fluorescent mScarlet protein fused to the localization signal peptide of dsbA, a periplasmic protein. The image obtained at t = 18 h shows co-localization of mScarlet (bottom, red) and pyoverdin fluorescence (top, blue, scale bar represents 10 *µ*m). B) Quantitative analysis of time-resolved images at 30 min and 18 h. Cells were segmented and intensity of fluorescence was measured along the long axis. Fluorescence at the poles was compared with the signal produced at the center of each cell; this was done for both pyoverdin (x-axis) and mScarlet (y-axis). Dots represent the intensity ratio for both fluorescent molecules. For measurements at 18 h the values for both the new pole (green dots) and old pole (yellow dots) are represented. Polar information is not available for freshly inoculated cells. The gray dashed line highlights the values where the center of the cell and the poles are not substantially different, i.e., their fluorescence ratio is close to 1. Deviation from this value indicates polar accumulation of the molecules. C) Cartoon depicting a simplified version of the pyoverdin pathway in which candidate genes for polarization are shown. 1. Pyoverdin (Pvd) synthesis starts in the periplasm in response to iron scarcity mediated by the transcription factor PvdS. Pyoverdin is then secreted to the bacterial periplasm, where it matures and becomes fluorescent. 2. Periplasmic pyoverdin is exported into the external medium by a complex that includes the transporter OpmQ. There, it chelates insoluble iron (Fe^3+^). 3. Ferripyoverdin complexes (no longer fluorescent) are then imported back into the periplasm after binding to the receptor FpvA. This receptor is known to also bind free pyoverdin [7]. In the periplasm, iron is extracted and pyoverdin is again recycled into the external medium by OpmQ. Polarization might also be determined by genes unrelated to the pyoverdin pathway: 4. SBW25 is known to secrete polymers such as cellulose that might trap pyoverdin [23]. 5. Pyoverdin could accumulate at the cell poles due to the rod shape of SBW25. D) Mutants associated to the main processes depicted in A) and their phenotype with regards to pyoverdin polarization. Mutants were grown on an agarose pad as described and fluorescence images displaying pyoverdin were taken at the time points indicated in the photo. The pyoverdin nonproducing mutant *pvdSG229A*(D77N) was co-inoculated with SBW25 to enable access of the mutant to pyoverdin. Mutants were tagged with a red fluorescent protein, one mutant colony is displayed in the image.

While overlap in patterns of pyoverdin and periplasm-targeted mScarlet accumulation points towards the fact that pyoverdin polarization is coupled to general cellular phenomena, possibilities remain for the involvement of genetic determinants of pyoverdin biosynthesis, regulation and / or transport. To test this hypothesis, patterns of pyoverdin accumulation were monitored in a range of mutants.

First, polar accumulation of pyoverdin is not a consequence of defective periplasmic maturation [25]: SBW25 producer and nonproducer types (the latter with an mCherry fluorescent marker) were co-cultured. Pyoverdin was localized in nonproducing cells (Fig. 3.1) demonstrating that accumulated pyoverdin is functional, since any siderophore internalized by Pvd^-^ necessarily comes from the external medium after export by SBW25. Polarization thus involves pyoverdin that is actively used among cells of the population.

Next, recognizing that mature pyoverdin interacts with membrane proteins via OpmQ [26, 27], *opmQ* was deleted and the pattern of pyoverdin accumulation determined. SBW25 Δ*opmQ* showed no change in capacity to accumulate pyoverdin at cell poles (Fig. 3.2), however, the mutant showed increased intracellular levels of pyoverdin; secretion was not abolished [28](Fig. S6 A).

Pyoverdin synthesis is subject to positive feedback control via the pyoverdin receptor FpvA making this protein a candidate for involvement in pyoverdin accumulation [7]. Production of pyoverdin by SBW5Δ*fpvA* was significantly reduced compared to the ancestral type (Fig. S6 B), however, where pyoverdin was visible (thus accumulated), it was typically polarized as in ancestral SBW25 (Fig. 3.3).

While the genes involved in extraction of ferric iron and translocation into the cytoplasm have not been characterized in *P. fluorescens*, a recent study in *P. aeruginosa* offered potential candidates for polarization including a siderophore-binding periplasmic protein [9]. Deletion of the homologs of these genes in SBW25 PFLU 2048, PFLU 2041, and PFLU 2043 did not disrupt accumulation of pyoverdin (Fig. S7). Thus, neither aggregation of pyoverdin at trafficking points across the periplasm, nor accumulation via the ferripyoverdin-complex recycling, are involved in polarization.

Additional possibilities for polar accumulation are via connections to extracellular polymer synthesis and cell morphology. SBW25 secretes cellulose when constructing bacterial mats at the air-liquid interface [23, 29] which could trap pyoverdin at the perimeter of cells. However, pyoverdin localization is not altered in SBW25 Δ*wsp*Δ*aws*Δ*mws* [30] that is unable to produce cellulose (Fig. 3.4). Finally, the rod-cell-shape characteristic of *Pseudomonas* was considered a possible contributory factor. To test this, pyoverdin accumulation was observed in a spherical Δ*mreB* mutant [31]. Despite the aberrant cell shape, this mutant displays fluorescence foci of pyoverdin after extended culture (Fig. 3.5). Curiously, pyoverdin accumulation at the pole is also independent of cell size in the ancestral genotype (Fig S9).

**Fig. 4.**
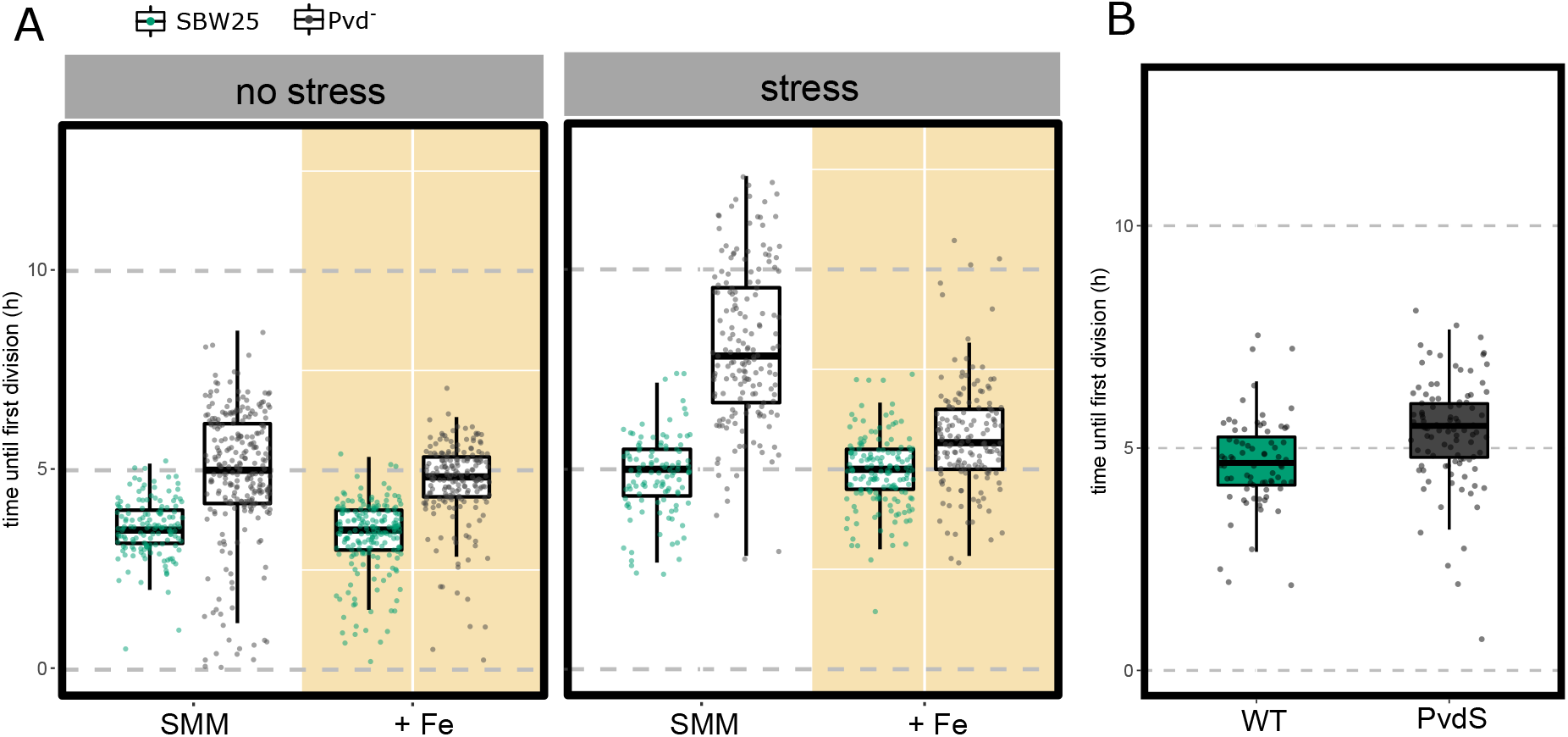
Pyoverdin accumulation facilitates recovery of growth after stress. A) Time until first division (lag time) of SBW25 (green) and pyoverdin defective mutant *pvdS229* (D77N), termed Pvd^-^ (gray) under different treatments and conditions. Prior to inoculation cells were either grown in the usual culture medium SMM (no stress on left panel) or treated with DP for 4h (stress on right panel). Cells were then inoculated on a fresh agarose pad, supplemented with 0.45 mM Fe_2_[SO_4_]_3_: these are labeled “+ Fe” (yellow background); unsupplemented SMM treatments are labeled “SMM” (white background). Dots represent individual cell values, box plots represent the associated distribution (median, 25th and 75h percentiles) N = 145, 261, 199, 193, 111, 168, 156, 156, respectively, from left to right. B) Time until first division of SBW25 (green) and Pvd^-^ mutants co-inoculated in a fresh agarose pad after separate stress treatment (4h in DP). Pvd^-^ mutants were labeled with a red fluorescent protein to allow identification of individual cells. Dots and box plots as in A). N = 175 total cells.

**Fig. 5.**
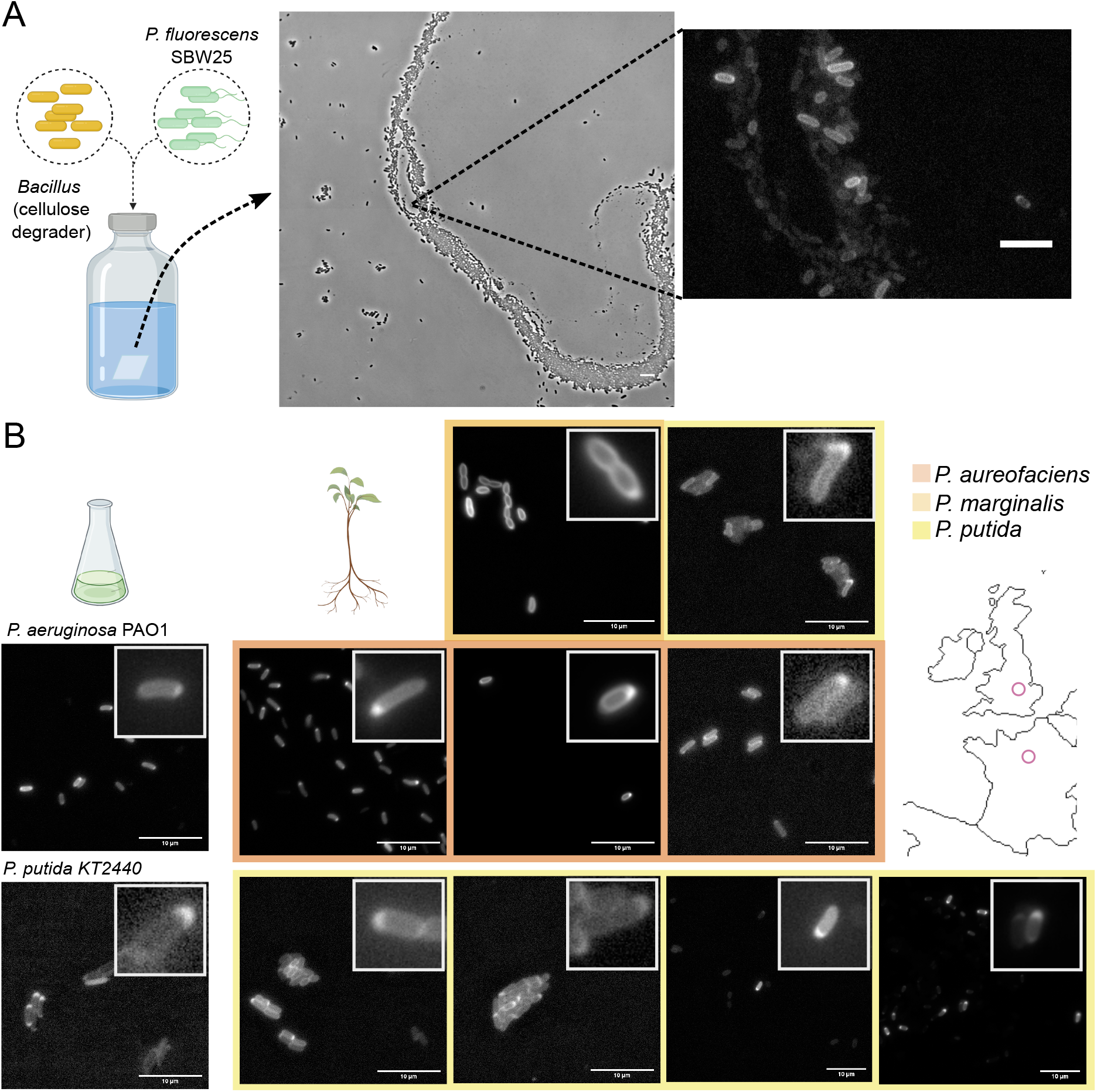
Accumulation visible by polarization is evident in SBW25 in a multispecies community and is a common phenotype in related species of the genus *Pseudomonas*. A) Polarized SBW25 cells in a multispecies community with interdependencies. SBW25 and a cellulose-degrading *Bacillus* strain isolated from a compost heap in Paris, France [32] were co-cultured in glass vials with minimal medium and cellulose paper as the only carbon source. The community was periodically sampled to assess the polarization state of SBW25. A representative image obtained after 28 days of growth is displayed, where both strains are visible (left, phase contrast image) and in a magnified region where pyoverdin distribution in SBW25 is visualized (right, fluorescence image). B) Qualitative accumulation assessment in species of *Pseudomonas* other than SBW25. A collection of laboratory (left) and natural (right) strains were tested for accumulation as evident by polarization. Accumulation by polarization in commonly used laboratory strains: *P. aeruginosa* PAO1 tested in SMM agarose pad with 1000 *µ*g/ml DP for 7:30 h; *P. putida* KT2440 tested in SMM agarose pad for 24h. Polarization test in natural isolates, left to right and top to bottom: U106 (liquid KB medium for 24 h); U177 (SMM agarose pad for 24 h); U149 (liquid SMM with DP 100 *µ*g/ml for 24 h); U180 (liquid SMM with DP 100 *µ*g/ml for 24 h); U181 (SMM agarose pad for 16 h); T24 V1a (SMM agarose pad for 24 h); T24 V9b (SMM agarose pad for 24 h); T24 H1b (SMM agarose pad for 24 h); T24 H9b (liquid SMM for 24 h). Natural isolates were collected in Oxford, UK [33] (first and second row) and in Paris, France [32] (Bottom row)(locations are roughly marked with a pink circle on the map). Color represents the identified *Pseudomonas* species. Insets highlight cells with clearly accumulated pyoverdin.

Polar accumulation of pyoverdin is linked to cell physiology, but is unperturbed by known genetic determinants of pyoverdin biosynthesis, regulation, or transport. This leads to the possibility that pyoverdin accumulation under conditions of cellular stress is an accidental consequence of the biophysics of cell biology and lacks ecological relevance. The alternate hypothesis is one of ecological relevance: a plausible explanation being that release of accumulated pyoverdin allows cells to reduce the time required to exit stationary phase.

An ideal test would be an experiment in which time to resumption of growth of an ancestral type that accumulates pyoverdin is compared to a mutant unable to accumulate pyoverdin, under conditions where pyoverdin accumulation both occurs, and doesn’t occur. If accumulation of pyoverdin confers a fitness advantage, then ancestral types are expected to resume growth more rapidly than non-accumulating mutants, but only after cells have first experienced conditions that promote accumulation of pyoverdin.

Unfortunately, as evident above, no such pyoverdin-accumulating mutant exists, nonetheless, progress is possible via an experiment that exploits the fact that a PvdS mutant cannot accumulate pyoverdin because it is unable to produce it. To this end, SBW25 and a PvdS mutant (SBW25 *pvdSG229A*(D77N) [13] (hereafter termed Pvd^-^) were grown separately in SMM for sufficient time to ensure cells entered stationary phase (24 h). A sample of cells from both populations was then transferred to independent SMM agarose pads and the time to first cell division determined. While a slight delay in median time for resumption of growth was observed in Pvd^-^ compared to SBW25 the distribution of data points shows significant overlap. The findings were not affected by transfer of cells from overnight SMM culture to agarose pads additionally supplemented with excess iron (Fig. 4A, left). This indicates that even the Pvd^-^ mutant was not deficient in iron despite having entered stationary phase.

Results were significantly different when the 24 h pre-culture period in SMM included a final 4 h period during which populations of both SBW25 and Pvd^-^ experienced iron-stress arising from supplementation of SMM with the iron chelating agent DP. As shown above, a 4 h treatment with DP not only caused cessation of growth, but also triggered polarization and accumulation of pyoverdin. The effect of this treatment was to greatly extend the time to resumption of growth in the non-pyoverdin-accumulating mutant. This difference was eliminated when cells, having experienced iron-deprivation, were provided with excess iron (Fig. 4A, right). These results demonstrate a fitness advantage to accumulation of pyoverdin that stems from a reduction in time to exit stationary phase.

This conclusion is further supported by the results of an additional experiment in which SBW25 and the Pvd^-^ mutant were grown as previously, treated with DP and then mixed in a 1:1 ratio and then transferred to a fresh SMM agarose pad. Measurement of the time to resume growth showed little difference among the competing genotypes (Fig. 4B). This is consistent with the expectation that SBW25 releases accumulated pyoverdin into the (fresh) medium to the benefit of both producer and non-producer alike. Notable though is the fact that producer cells reap greatest benefit. When nonproducers are instead inoculated in the presence of rare ancestor cells (∼ 1%) their lag time again increases significantly, supporting the conclusion that newly released pyoverdin underpins the faster division of producers (Fig. S10). Furthermore, the amount of pyoverdin accumulated during starvation seems to be sufficient to support at least twice as many cells, as the lag time of SBW25 producer cells was not affected by the presence of nonproducers.

We further tested pyoverdin polarization in conditions closer to the natural milieu of SBW25, where species inter-dependencies are common and resources invariably limiting. To implement a minimal bacterial community, SBW25 was co-cultured with a cellulose-degrading *Bacillus* isolated from a compost heap [32]. Because SBW25 is unable to degrade this polymer, it must rely on the *Bacillus* species to obtain carbon for growth. These two strains were grown together in minimal medium with cellulose paper as the sole carbon source and periodically imaged to assess polarization status. Pyoverdin was readily observed to be polarized (Fig. 5 A). Pyoverdin accumulation at the cell pole is thus likely to be part of the natural phenotypic repertoire of SBW25, and given that in natural environments bacteria likely find themselves in or close to stationary phase most of the time, polar accumulation of pyoverdin stands to be the normal state.

Since pyoverdins are produced by many species of the genus *Pseudomonas*, investigating subcellular distribution in related strains may provide an evolutionary context for polar accumulation. We selected 11 strains belonging to the genus *Pseudomonas*, comprising both common laboratory strains (such as *P. aeruginosa* PAO1, in which pyoverdin is usually studied) and natural isolates from different European locations. The ability of these strains to polarize pyoverdin was classified qualitatively, that is, presence or absence of polarized cells (Fig. 5 B.) All tested strains localized pyoverdin at the cell poles in conditions similar to those described here for SBW25. Interestingly, different strains polarized pyoverdin with treatments of varying stringency. In some cases extended culture in SMM was enough to observe the phenotype (e.g., *P. putida* KT2440). In others, amendment with an iron chelator was necessary to observe the effect (e.g., *P. aureofaciens* U149), at the same dosage that induced polarization in SBW25, and in the specific case of *P. aeruginosa* PAO1, very high doses of DP were required. This range of responses could reflect the secretion of secondary siderophores by some strains [34] or differences in the regulation of pyoverdin production [12], and suggest that polarization is an ecologically relevant trait that varies depending on the evolutionary history of the lineage.

## Discussion

Previous work showing that the population-level distribution of pyoverdin changes depending on nutrient status [13], contact with neighboring cells [14], and environmental stress [15] motivated our investigation. With focus on *P. fluorescens* SBW25, and using time-resolved microscopy, we have shown that pyoverdin transiently accumulates within cells (preferentially at cell poles), that localization is a reversible process associated with arrest of cell division, and is affected by factors such as entry into stationary phase and deprivation of specific nutrients (Fig. 2). Particularly significant is demonstration that accumulation of pyoverdin has ecological relevance (Fig. 4).

Investigations at the level of individual cells allow cell-level behaviors to be linked to the dynamics of populations [17, 18]. For example, Gore et al (2009) [35] showed that the cellular location of invertase responsible for degradation of sucrose in *Saccharomyces cerevisiae* creates diffusion gradients of degradation products that deliver benefit to producing cells, despite costs to producers of synthesizing the enzyme [35]. Similarly, the siderophore enterochelin (a catecholate secreted by *E. coli* and other *Enterobacteriaceae*) can remain associated with the outer membrane under conditions of low cell density, delivering preferential benefit to enterochelin-producing cells [36]. Some localized products display dynamic behaviour [37], with, in some instances, proteins oscillating between poles [38]. In other instances, proteins accumulate via formation of nuclear occlusions in the cytoplasm [39]. In the context of pyoverdin, complexes termed “siderosomes” localize enzymes for synthesis of pyoverdin at old cell poles during exponential growth by association of the cytoplasmic membrane with L-ornithine N5-oxygenase (encoded by *pvdA*) [40, 41].

While pyoverdin in SBW25 was most often observed at new cell poles, it was also evident at old poles; on occasion it was found at both poles. Moreover, although all cells accumulate pyoverdin (Fig 1C), a minority show no evidence of polar localization. Measurement of the amount of pyoverdin in cells that localize the iron chelator showed that it is the same as that contained within cells displaying a homogeneous distribution of pyoverdin (Fig S2).

Taken together, ambiguity surrounding accumulation and polar localization suggested a general connection to biophysical aspects of cell biology. Interestingly, a recent study of *E. coli* under starvation conditions showed polar accumulation of fluorescent markers, including mCherry [24]. Accumulation was connected to shrinkage of the cytoplasm, with shrinkage creating an expansion of space in the polar region of the periplasm. It thus seemed plausible that accumulation of pyoverdin in SBW25 in stationary phase may be a consequence of nonspecific cellular behavior under growth arrest. Such a possibility was bolstered by the fact that a PvdS mutant deficient in synthesis of pyoverdin (and thus lacking siderosomes) nonetheless accumulated pyoverdin, that deletion of genetic determinants of uptake and recycling had no observable effects on accumulation, and that pyoverdin accumulation also occurred at discrete locations in spherical Δ*mreB* cells (Fig. 3D.5).

To investigate the hypothesis that pyoverdin accumulation is connected to cytoplasmic shrinkage, red fluorescent mScarlet was translationally fused to the localization signal peptide of DsbA. As cells entered stationary phase accumulation of the fluorescent marker protein was observed at cell poles in a manner analogous to that observed for pyoverdin. In fact, single images captured using filters specific for the reporter protein and pyoverdin showed an almost perfect overlap (3A,B). This lead us to conclude that polar accumulation of pyoverdin is part of a general cellular response to starvation. Thus observations made in *E. coli* with a chimeric reporter [24] appear to hold for SBW25, but with our observations connecting cytoplasmic shrinkage to accumulation of a biologically relevant molecule. Additionally we demonstrate that accumulation delivers beneficial effects on fitness.

Data in Fig. 4A show that cells that have accumulated pyoverdin – most of which is localized to cell poles – resume growth more rapidly compared to cells that have not accumulated pyoverdin. A time-to-first cell division advantage was not evident when growth-arrested cells were transferred to iron-replete conditions. This demonstrates that liberation of the stock of pyoverdin accumulated during growth cessation speeds the time to growth resumption after growth arrest, presumably through provision of available iron to growing cells 4B.

Of additional interest is evidence from cell-level observations that rare pyoverdin producers are largely unaffected by the presence of Pvd^-^ mutants (Fig. S10). Such mutants have often been referred to as “cheats” and thus expected to negatively impact the fitness of pyoverdin producers (so called “cooperators”). Lack of detrimental impact is further evidence that pyoverdin production preferentially benefits producer cells. This finding further calls into question the fit between social evolution theory and production of extracellular products by microbes [13, 12]. It also gels with work showing that populations of siderophore-producing *Pseudomonas* can be grown in the presence of an excess of nonproducing mutants without significant reduction in overall yield [42].

Recent work suggests important physiological differences between acute (and unexpected) interruptions of cell division, and gradual growth arrest that determines entry into stationary phase [**?**]. In the former case, sudden arrest induces a disrupted state that results in large variability in cell-level duration of lag phase. In the latter case, cells implement genetic programs that buffer the effects of impending starvation creating cohesive population-level responses [**?**]. The seemingly adaptive nature of pyoverdin accumulation under stress is reminiscent of – and perhaps even connected to – the capacity of SBW25 to enter a semi-quiescent capsulated state upon starvation. During starvation, SBW25 cells produce an excess of ribosomes that allow rapid exit from stationary phase once growth-permissive conditions are encountered. Cells unable to provision ribosomes are quickly out-competed by those that do [43]. It is possible that localization of pyoverdin, followed by fast release under growth permissive conditions, has evolved as a strategy precisely because it maximizes competitive performance upon resumption of growth. Rapid resumption of growth is likely to deliver significant fitness benefits in environments punctuated by periods of nutrient abundance and scarcity.

Mounting evidence indicates that pyoverdin – in its many variant forms [34] – has multiple impacts on composition and function of microbial communities [44, 45, 46, 47, 48, 49]. The fact that pyoverdin is accumulated and localized under growth limiting conditions in a diverse range of *Pseudomonas*, combined with evidence of the same behavior in SBW25 cells grown for weeks under nutrient restrictive conditions with reliance on cellulose degrading *Bacillus* (Fig. 5), suggests that polarization has relevance to conditions beyond those experienced by cells in standard, nutritionally-rich, exponential-phase, laboratory culture.

Given that pyoverdin accumulation appears connected to cytoplasmic shrinkage, it seems reasonable to assume that other molecules may similarly accumulate in the periplasm in response to division arrest signals, which both change size and shape of cellular compartments. Indeed, accumulation of products related to nutrient stress is one possible function of the bacterial periplasm, with roles analogous to vacuoles in eukaryotes [50].

## Materials and methods

### Strains, culture media, and reagents

Ancestral *Pseudomonas fluorescens* SBW25 originally isolated on beet roots at the University of Oxford farm (Wytham, Oxford, U.K.) [20] and a collection of relevant mutants were used: the pyoverdin nonproducer *pvdsG229A*(D77N) named Pvd^-^ in the main text (construction described in [13]), the corresponding strain with mCherry fluorescence tagging under IPTG induction, Δ*mreB*, PBR716 (Δ*wsp*Δ*aws*Δ*mws*, described in [30]). Pyoverdin import and export defective mutants Δ*PFLU3979* (OpmQ) and Δ*PFLU2545* (FpvA) were created by two-step allelic exchange [51]. The periplasm reporter strain was created by fusing the red fluorescent protein mScarlet to the signal peptide of periplasmic protein, DsbA (see Supplementary Methods). A neutrally marked SBW25 strain [52] was used to replicate the fitness assays from [13]. *Escherichia coli* DH5*α λ*_pir_ and pRK2013 were used for cloning. *Bacillus* 002.IH from Steven Quistad’s compost heap collection [32] was used for the community experiment. For the phylogenetic comparison, an assortment of species of the genus *Pseudomonas* were selected from the lab collection. Most experiments were performed in succinate minimal medium (SMM, described in [14]). Overnight cultures were done in Luria-Bertani (LB) broth. Replication of fitness assays was performed in CAA and KB media as described in the original work [13]. Community experiment was done in M9 minimal medium with cellulose as the only carbon source (Whatman). Where indicated media was supplemented with 2,2’-dipyridil (Sigma), tetracycline (Duchefa, France), Fe_2_[SO_4_]_3_(III) (Sigma), IPTG (Melford). For strain construction tetracycline, nitrofurantoin (Sigma), X-gal (Melford), D-cycloserine (Duchefa) were used.

### Agarose pad

To prepare the agarose pad, 220 *µ*L of agarose (Melford) dissolved in SMM (2% w/v) were poured onto a microscope slide fitted with a sticky frame (Gene Frame, Fisher Scientific), pressed with a clean slide and allowed to dry for ∼ 2 min. A small ∼ 3 mm section of the pad and frame was cut across the slide to provide air for the growing cells. 1.5 *µ*L of washed culture was inoculated on the agarose pad and sealed with a coverslip.

### Microscopy

Inoculated agarose pads were monitored by taking snapshots every 30 min for a typical total time of 18h using the microscope Axio Observer.Z1 (Zeiss, Germany). Cells were imaged under phase contrast (exposure: 100 ms) and fluorescence corresponding to Pvd using the fluorescence Source X-Cite 120 LED and the following filters: 390/40 BrightLine HC, Beamsplitter T 425 LPXR, 475/50 BrightLine HC (exposure: 30 ms, 12 % intensity). Images were taken with 63x and optovar 1.6x magnification.

### Image analysis

Image processing was performed using the image analysis software Image J. Segmentation and analysis was performed using the package for Matlab SuperSegger from the Wiggins Lab [53]. Further analysis and data visualization was carried out with the programming software R. After segmentation cells were sorted by polarization status using a classificator obtained with the Statistics and Machine Learning Toolbox for Matlab (More information in Supplementary Methods).

### Community experiment

SBW25 and *Bacillus* 002.IH (from [32]) were grown on 20 ml M9 minimal medium and cellulose as the only carbon source (1 cm × 1 cm cellulose paper). After overnight growth cultures were washed and co-inoculated into 20 ml of M9 medium with cellulose paper in a 60 ml vial. These vials were incubated at room temperature with unscrewed caps to allow air exchange, and periodically sampled for imaging.

### Phylogenetic comparison

A collection of laboratory and natural strains belonging to the genus *Pseudomonas* were subjected to a binary qualitative polarization test, i.e. the test was considered positive if polarized cells were observed under fluorescence microscopy in conditions similar to SBW25 but no dynamics were assessed. Commonly used laboratory strains *P. aeruginosa* PAO1 and *P. putida* KT2440 were tested. Natural isolates belong to collections from two different locations. Paris, France [32]: T24 V1a, T24 V9b, T24 H1b, T24H9b; all classified as *P. putida*. Oxford, UK [33]: *P. marginalis* U106, *P. putida* U177, *P*.*aureofaciens* U149, U180, and U181.

## Author contributions

PBR, CMF and MA designed research. CMF performed research. CMF and MA analyzed data. PBR, CMF and MA wrote the paper.

## Acknowledgements

We thank Dave Rogers and Ellen McConnell for assistance with construction of mutants, Steven D. Quistad and Xue-Xian Zhang for natural strain collections, and Petra Levin and Adrienne Brauer for useful discussion. PBR acknowledges generous core support from the Max Planck Society.

## Supplementary Appendix

### Supplementary Methods: Image processing and phenotypic classification of cells

Images were pre-processed for segmentation using the image analysis software ImageJ [54]. Time-lapses of growing microcolonies were segmented using the SuperSegger Toolbox [53] for Matlab [55]. This toolbox has been designed specifically for the analysis of bacterial microcolonies imaged under phase contrast and fluorescence microscopy and allows the user to adapt segmentation variables to suit a particular experimental system using neural network training. Individual images of cells and their lifetime properties (including time of birth and death, lineage, polar identity, and others) are the ouptut of the software. This enabled tracking of cells throughout three generations mentioned in the main text: F0 or inoculum cells, F1 or exponentially growing cells resulting from the division of F0, and F2 cells or stationary phase cells resulting from the division of F1. Pyoverdin polarization is thus observed in F2 cells. To obtain a means to automatically classify cells by their pyoverdin polarization status the Statistics and Machine Learning Toolbox for Matlab was used [56]. 646 segmented cell images were collected as the training set, originating from 4 different experiments. The training set included images obtained from microcolonies growing on an agarose pad and from cells treated with an iron chelator in liquid medium, to account for differences between experimental setups that could impact on image analysis (e.g., overlapping cells in microcolonies, changing levels of relative cell fluorescence in different environments, etc.) These cells were visually classified into “polarized” and “homogeneous”. Next, image features were extracted to develop a classification algorithm. To extract uni-dimensional features from 2-dimensional images, these were processed as follows: first, mean fluorescence along the long cell axis was obtained (also named kymograph). Cells were then divided into three equally sized regions representing the center of the cell and both cell poles and different variables were obtained comparing these regions. Many variables were tested and optimum results were obtained when using the following procedure: kymographs were normalized by either subtracting or dividing by the median fluorescence of the cell center. From these, standard deviation, and 75th percentile of either polar region were calculated. These 6 features were fed into the Classification Learner App in Matlab with 5-fold cross-validation. The model with best results was a linear SVM (Support Vector Machine) model, with accuracy *>*90%. It is estimated that the model has a false negative rate of ∼5% for “homogeneous” cells and ∼10% for “polarized” cells i.e. it has a slight bias towards under-identification of polarization which we found more desirable than over-identification. This model was used for all subsequent classification mentioned in the main text and SI. To further analyze and visualize data the programming software R was used [57].

### Supplementary Methods: Construction of strain with periplasmic red fluorescent marker

The IPTG-inducible periplasmic mScarlet reporter was created by first introducing the mScarlet-I CDS from plasmid pMRE-Tn*7* -155 (amplified using primers mScarlet fwd and mScarlet rev) into plasmid pUC18-mini-Tn*7* T-LAC (linearized using primers Tn*7* LAC fwd and Tn*7* LA rev) using NEBuilder HIFI DNA Assembly Master Mix (New England BioLabs)[58, 59]. The DsbA signal sequence from SBW25 (the first 81 nucleotides of PFLU0083) was then introduced to the start of mScarlet-I in the resulting plasmid by FastCloning [60] using primers dsbAmScarlet fwd and dsbAmScarlet rev. The forward primer included the LEGPAGL amino acid linker used by Uehara *et al* (2009) [61] to construct dsbA_ss_-mCherry in *E. coli*. This final plasmid was introduced into SBW25 by electroporation (mediated by the transposition helper plasmid pUX-BF13 [62]) and integration of the transposable element containing Plac-dsbA_ss_-mScarlet-I into the attTn*7* site downstream of *glmS* was selected by plating to gentamicin. Correct integration of the Tn*7* cassette was confirmed by Sanger sequencing. All cloning was performed in *E. coli* OneShot TOP10 chemically competent cells (ThermoFisher Scientific) and PCRs were conducted using either Phusion High-Fidelity PCR Master Mix (ThermoFisher Scientific) or Q5 High-Fidelity 2X Master Mix (New England BioLabs). pMRE-Tn*7* -155 was a gift from Mitja Remus-Emsermann (Addgene plasmid # 118569 ;http://n2t.net/addgene:118569; RRID:Addgene_118569). pUC18-mini-Tn*7* T-LAC was a gift from Herbert Schweizer (Addgene plasmid # 64965 ;http://n2t.net/addgene:64965;RRID:Addgene_64965).

Primers (overlaps for assembly or primer extensions in lower case, annealing region in uppercase, dsbAss in italics, LEGPAGL linker in bold)

**Table 1.**
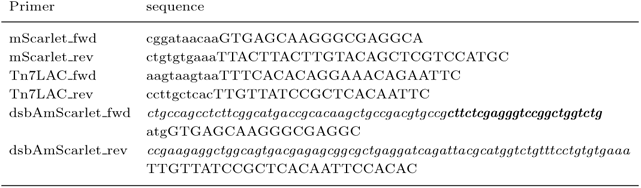
gasajcgaj.

**Fig. S1.**
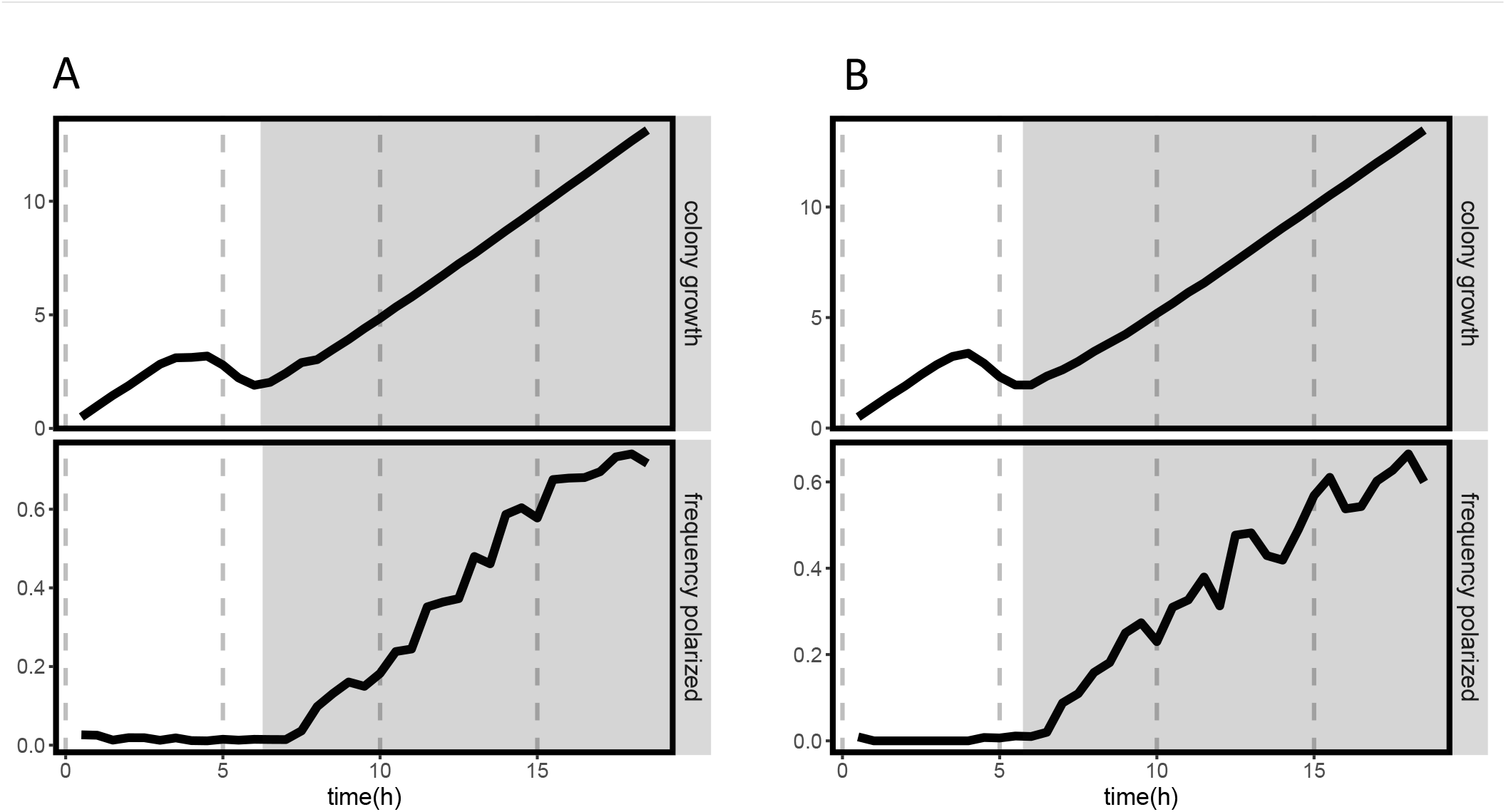
SBW25 consistently polarizes pyoverdin upon entry into stationary phase. Polarization in different growth stages of a microcolony. Both plots represent mean age of the cells in a growing microcolony of SBW25 (black line, top) and the corresponding frequency of polarized cells for each time point (black line, bottom). Bacterial colonies undergo different growth stages, from left to right: lag phase (generation F0), exponential phase (generation F1), and stationary phase (generation F2) Colored panels highlight stationary phase of bacterial colony growth, where pyoverdin is observed to accumulate at cell poles. In all cases data has been filtered to exclude cells with segmentation errors or other artifacts that preclude proper analysis. Each plot depicts a biological replicate of the experiment described in Fig. 2A. A) 5 technical replicates, N = 152 initial cells. B) 4 technical replicates, N = 109 initial cells

**Fig. S2.**
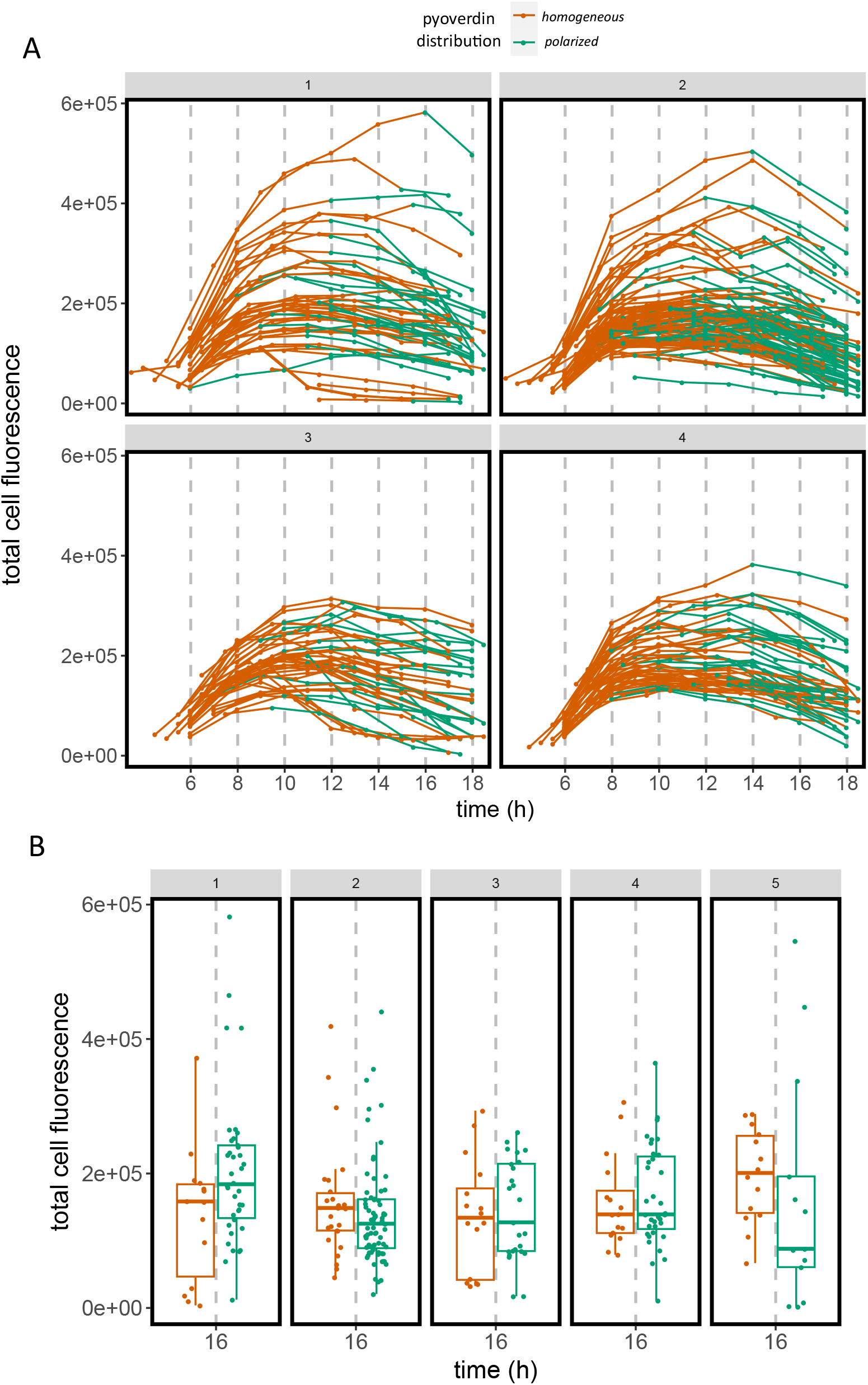
Total cell fluorescence over time in cells that accumulate pyoverdin at the pole during stationary phase. A) Each line represents an individual cell that is generated by cell division but does not subsequently divide over the course of the time-lapse, i.e. generation F2 as defined in the main text. Values are the sum of the fluorescence intensity of all pixels corresponding to a segmented cell. Lines are colored according to the pyoverdin distribution inside the cell at each time point, classified by a machine learning algorithm in “homogeneous” (red lines) or “polarized”(green line). Each panel labelled 1-4 represents a technical replicate, i.e. a different position in the microscope slide during image acquisition. B) Distribution of total cell fluorescence in cells that are classified as “homogeneous” (red) or “polarized” (green) at a time point corresponding to mid to late stationary phase, t = 16h. Box plots represent the mean and associated statistical parameters. Dots represent individual cell values. Panels represent technical replicates as in A.

**Fig. S3.**
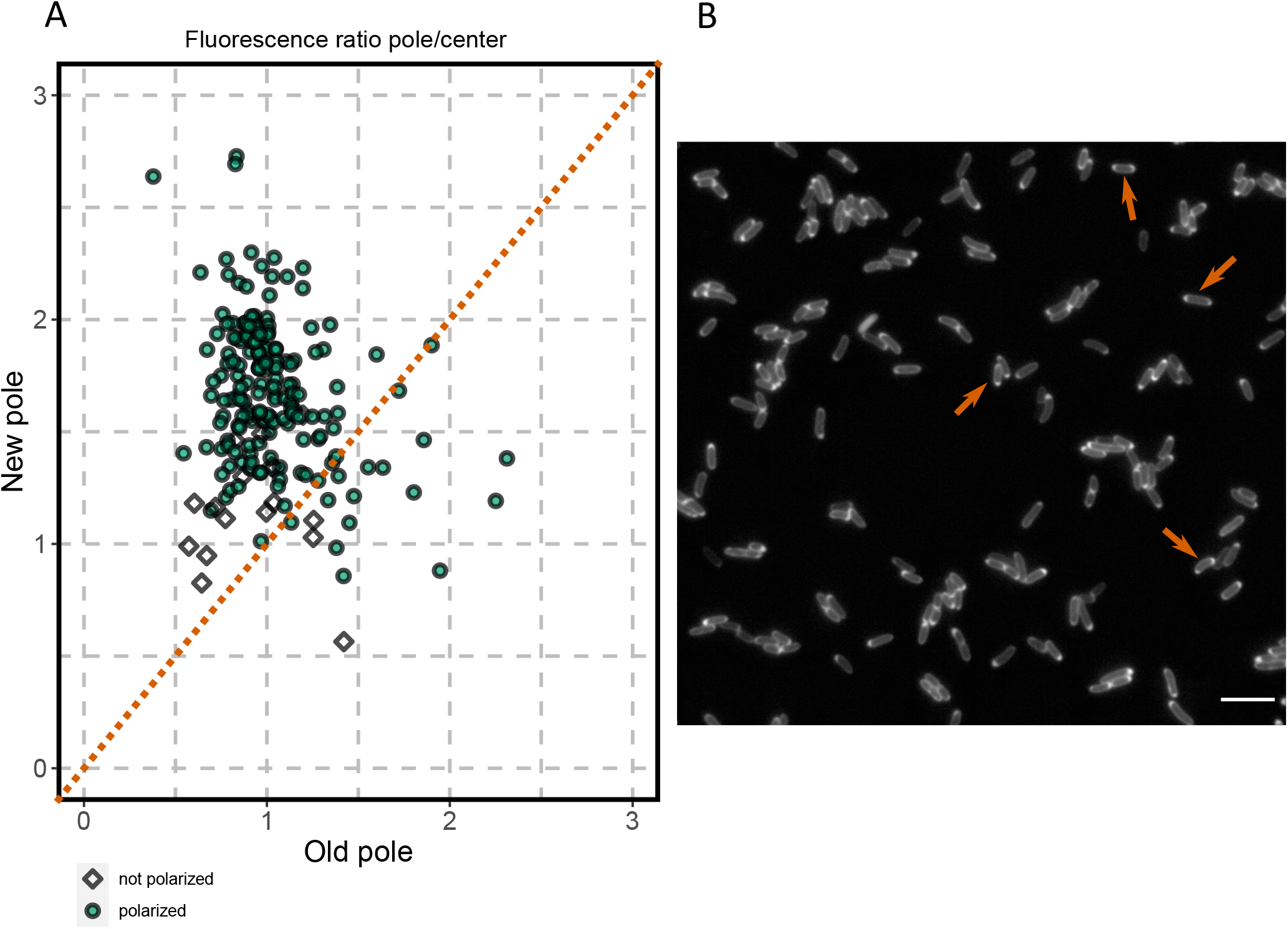
Accumulation of pyoverdin occurs predominantly at the new cell pole, but not exclusively and occasionally at both cell poles. A) Ratio of fluorescence between the central region of cells and the region of the new cell pole or old cell pole. Fluorescence values are obtained by dividing segmented cells into 3 proportional regions over long cell axis and averaging fluorescence along both cellular dimensions. Data corresponds to individual F2 cells at time = 18 h, N = 187. Green circles represent cells classified as “polarized”, white diamonds represent cells classified as “not polarized”. Red line highlights 1:1 ratio of fluorescence in cell centers to cell poles for reference. B) Image of SBW25 cells in conditions that promote polarization, in this case, high cellular density leading to rapid entry into stationary phase (t = 18 h) where accumulation of pyoverdin at both cell poles can be observed in few individuals (red arrows). Scale bar = 10 *µ*m.

**Fig. S4.**
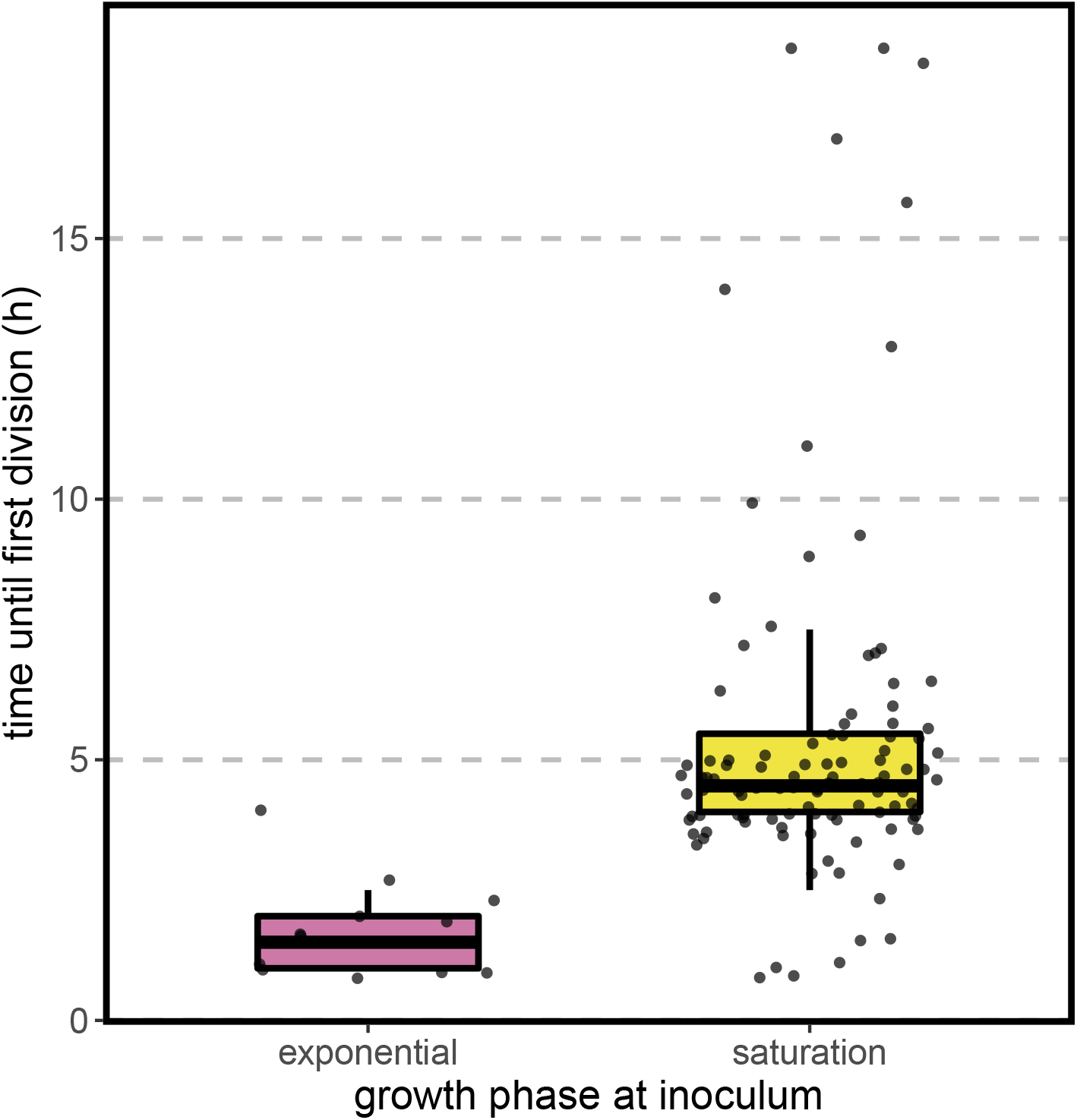
Lag time difference between cells in stationary and exponential phase. To assess the physiological state of cells after 24 h in SMM (whether they are still in exponential phase due to the minimality of the medium or rather they are entering stationary phase) we compared the time until first division of cells previously resuspended in fresh liquid SMM culture for 4h with data from cells inoculated in a fresh agarose pad directly following the usual procotol described in the main text. In the former case the cells are in exponential phase as their lag time is comparable to division time in this medium. (N = 13, 109 respectively)

**Fig. S5.**
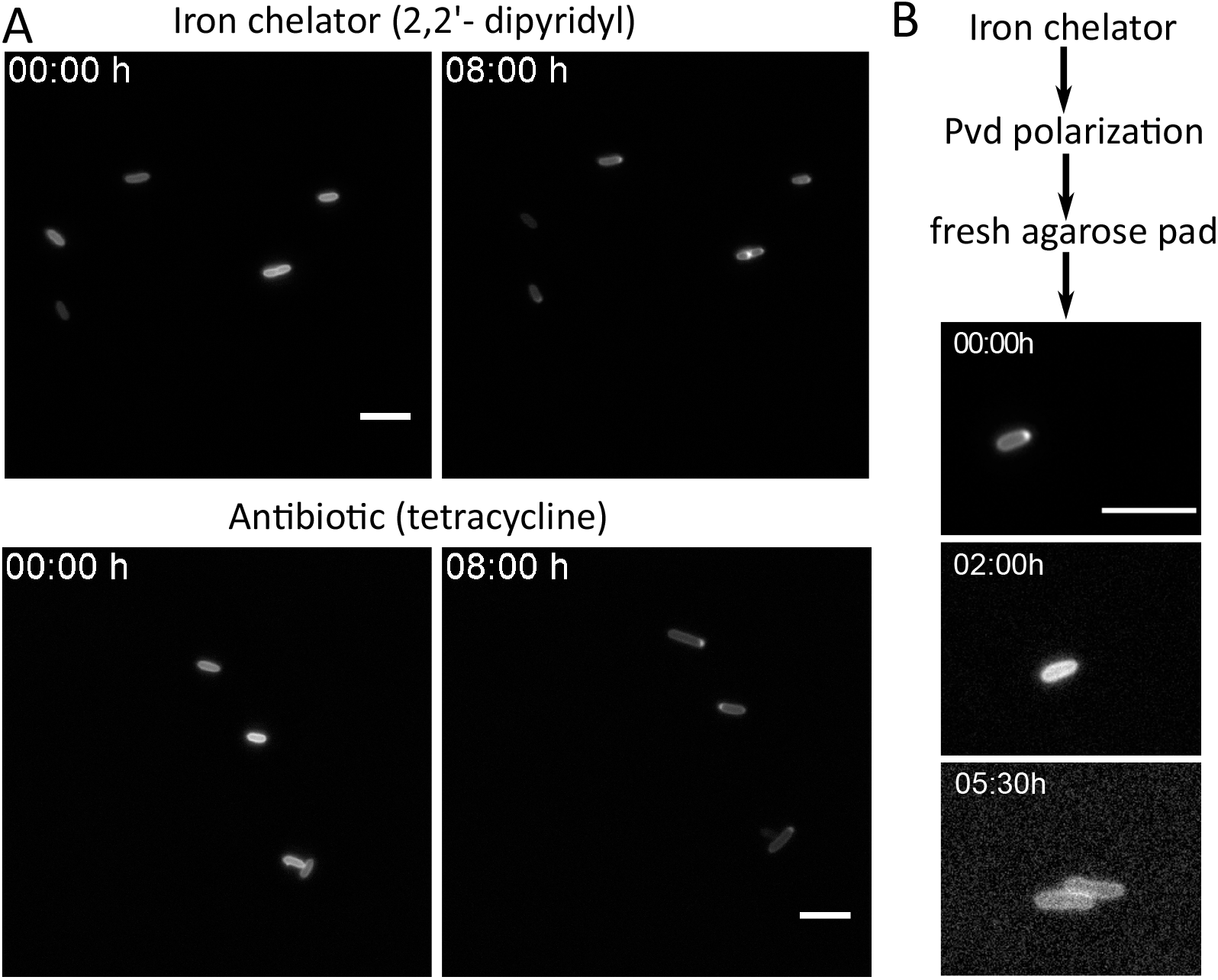
Exposure to chemicals that prevent cell growth induces reversible accumulation of pyoverdin at the cell pole. A)Treatment with iron chelator (top) or antibiotic (bottom) triggers accumulation of pyoverdin at the cell pole without detectable cell growth. SBW25 cells were inoculated on an agarose pad containing either 100 *µ*g/ml of 2,2’-dipyridyl (top) or 5 *µ*g/ml tetracycline (bottom). Images were taken at t = 0 h and t = 8 h at the same position on the agarose pad. Note then that the photos depict the same individual cells at both time points, as no cell division was observed. B) Pyoverdin polarization is reversible and precedes recovery of growth after stress. Time-lapse images of SBW25 pre-treated with an iron chelator to induce pyoverdin polarization, then washed and inoculated on a fresh agarose pad. Images follow one individual cell as it recovers homogeneous pyoverdin distribution in the periplasm, followed by cell division.

**Fig. S6.**
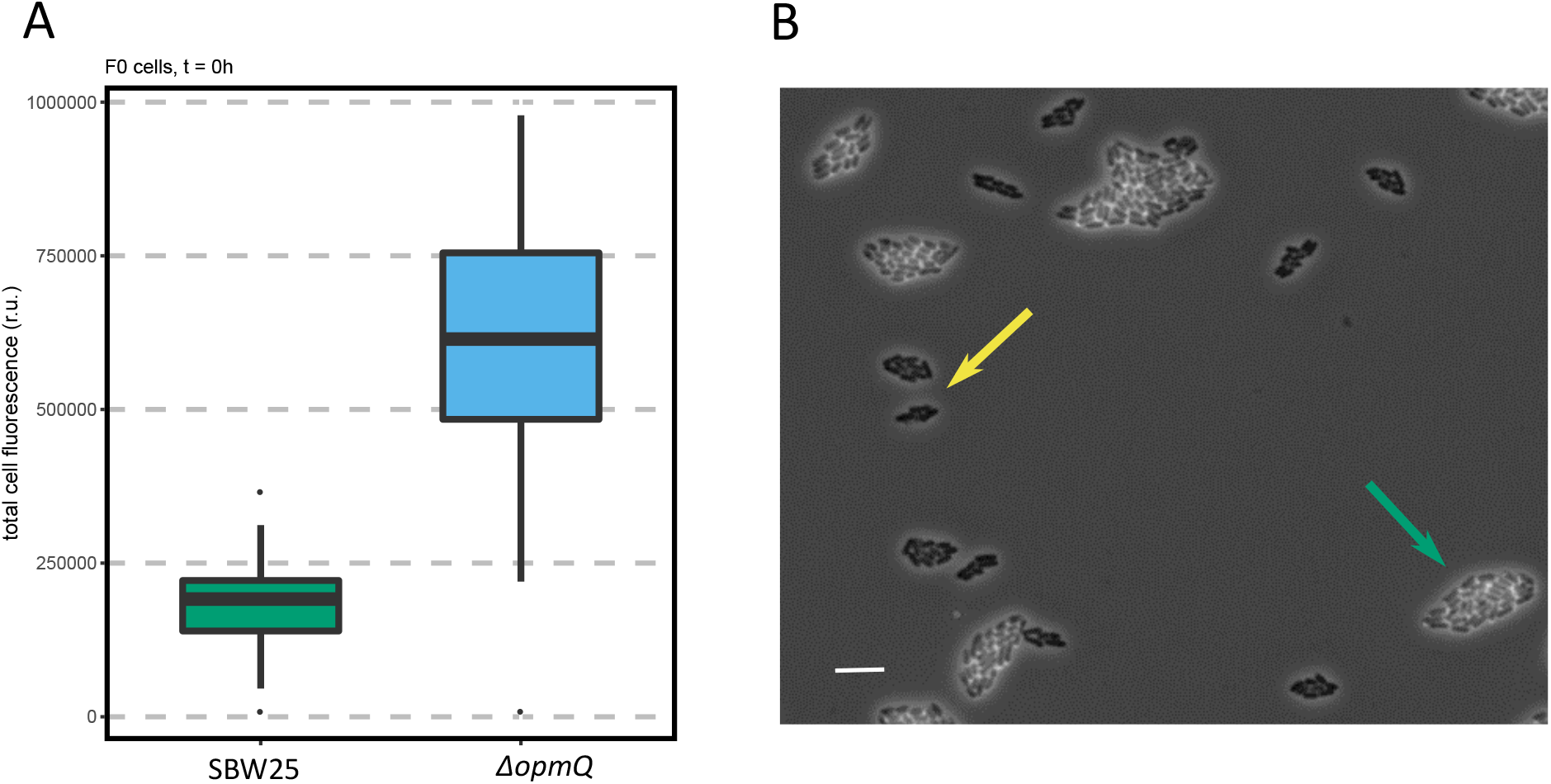
OpmQ and FpvA deletion mutants are impacted on their ability to recycle, import, and synthesize pyoverdin. A) Deletion of the export pump OpmQ leads to increased intracellular levels of pyoverdin compared to ancestral SBW25. Box plots represent the aggregated fluorescence of individual cells of SBW25 (green) and Δ*opmQ* (light blue) at t = 0 h of inoculation on the agarose pad, N= 35, 29, respectively. B) Deletion of the receptor FpvA reduces pyoverdin synthesis to basal levels, and impedes its import by crossfeeding. Image depicts co-culture of ancestral SBW25 (tagged with mCherry for identification) and Δ*fpvA*, where phase contrast and fluorescent imaging of pyoverdin have been overlayed to highlight the differences in pyoverdin production between both strains. Scale bar = 10 *µ*m. Strains were mixed in equal proportions and grown on an agarose pad for 18 h. Green arrows point to microcolonies of SBW25, yellow arrows point to Δ*fpvA* microcolonies, as representative examples of the two phenotypes.

**Fig. S7.**
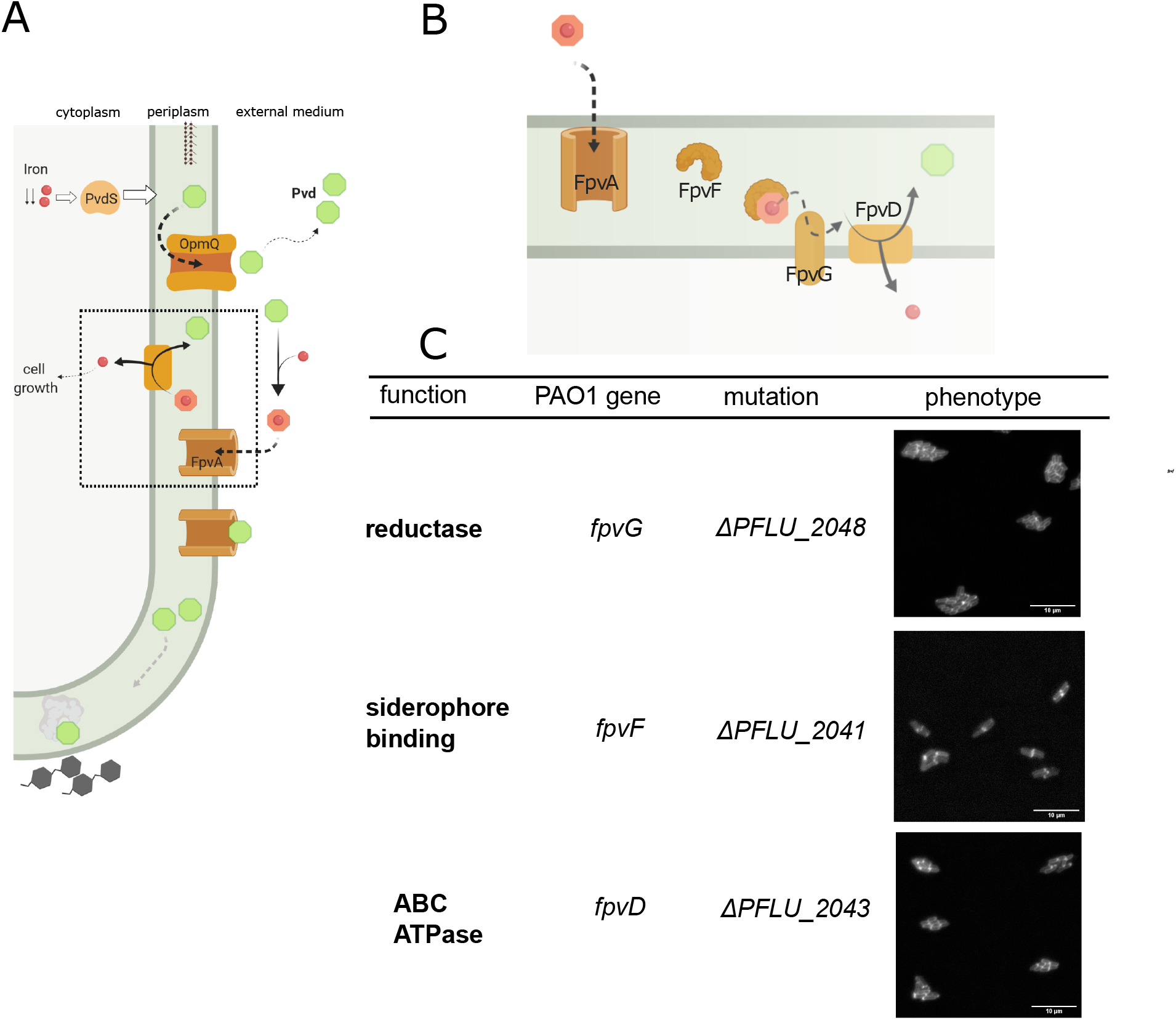
title. A) Cartoon depicting the pyoverdin pathway, see main text for details. B) Cartoon detailing the mechanism of iron extraction from the ferripyoverdin complex and recycling of pyoverdin molecules. C) Pyoverdin polarization phenotype of a collection of deletion mutants of the processes depicted in B). Genes were selected based on recent work on *P. aeruginosa* PAO1 identifying the function of previously unknown proteins. Orthologs of these genes were identified in *P. fluorescens* SBW25 in the database pseudomonas.com. Genes were deleted by two-step allelic exchange and cells were cultured and imaged as previously described.

**Fig. S8.**
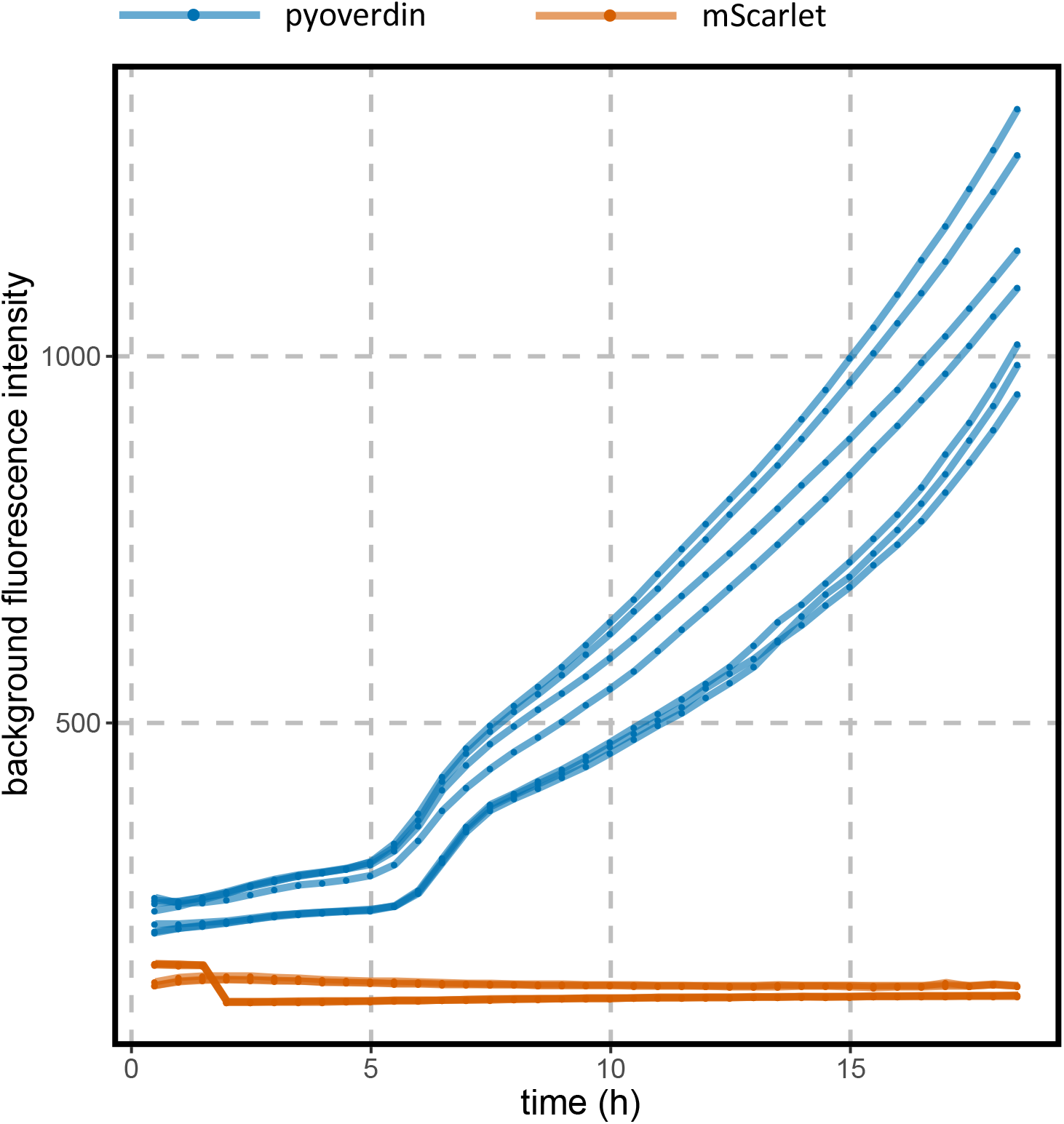
Fluorescence background measured when imaging cells that produce the naturally-occurring molecule pyoverdin and an engineered periplasmic mCherry protein. Dots and lines represent the median intensity value of a 100×100 px square in the microscope field where cells are not present. Red lines correspond to the red fluorescent protein mCherry while blue lines correspond to pyoverdin (which exhibits a fluorescence emission peak at 450nm). Each line represents a technical replicate.

**Fig. S9.**
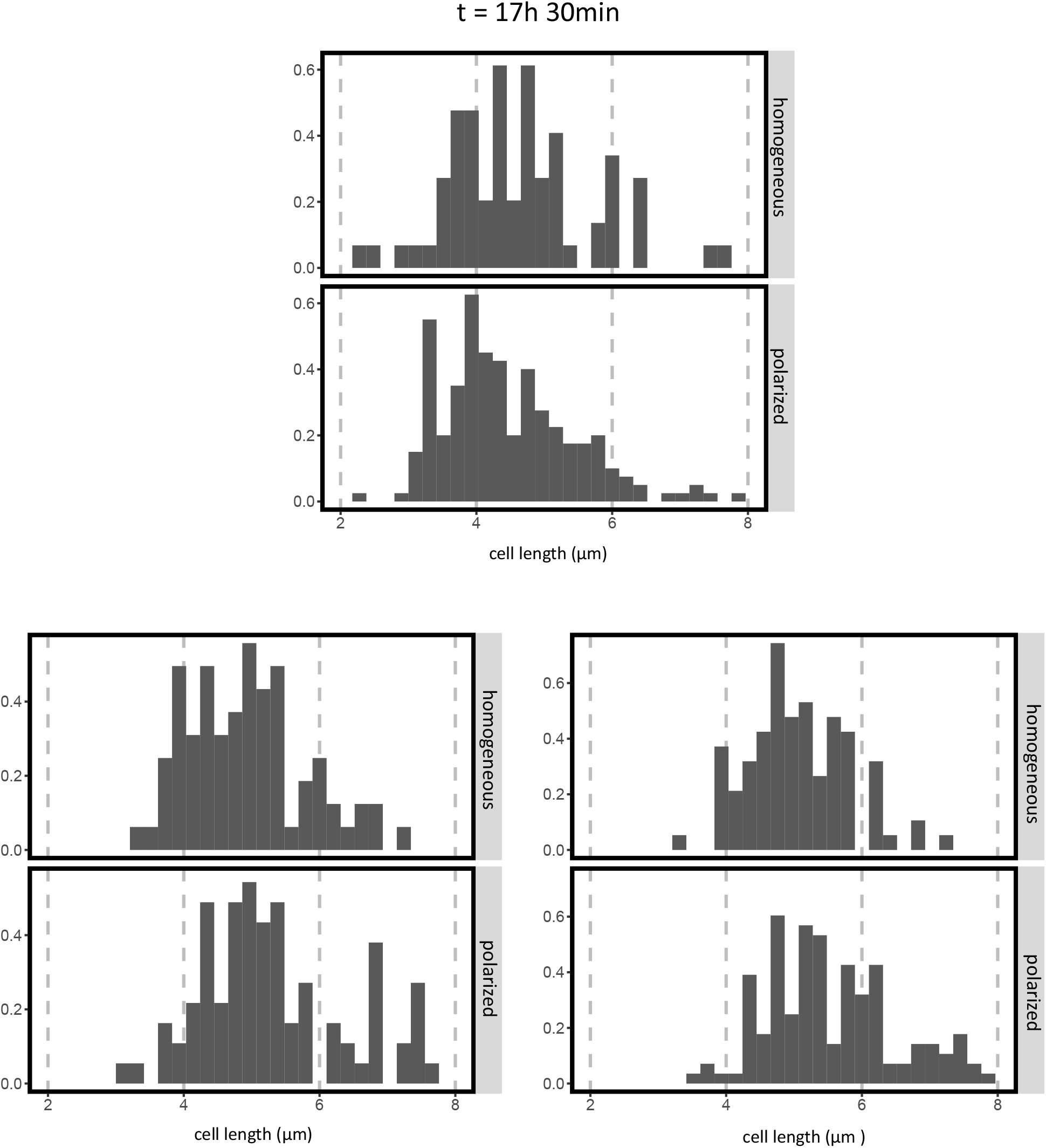
Accumulation of pyoverdin as a function of cell size. Distribution of cell size after 17h 30 min of growth in the agarose pad. The population of cells is classified according to the localization of pyoverdin in “accumulated”and “non-accumulated”. Cell size is measured as length on the long axis, since elongation occurs in this direction and thus will determine final cell area. Bars represent the distribution of cell lengths in both groups of cells. Each plot represents an independent experiment. Strains used: top: SBW25, bottom: MPB25340 (SBW25 with periplasmic red fluorescent marker), left and right panels represent 2 biological replicates. N = 261, 167, 235 respectively.

**Fig. S10.**
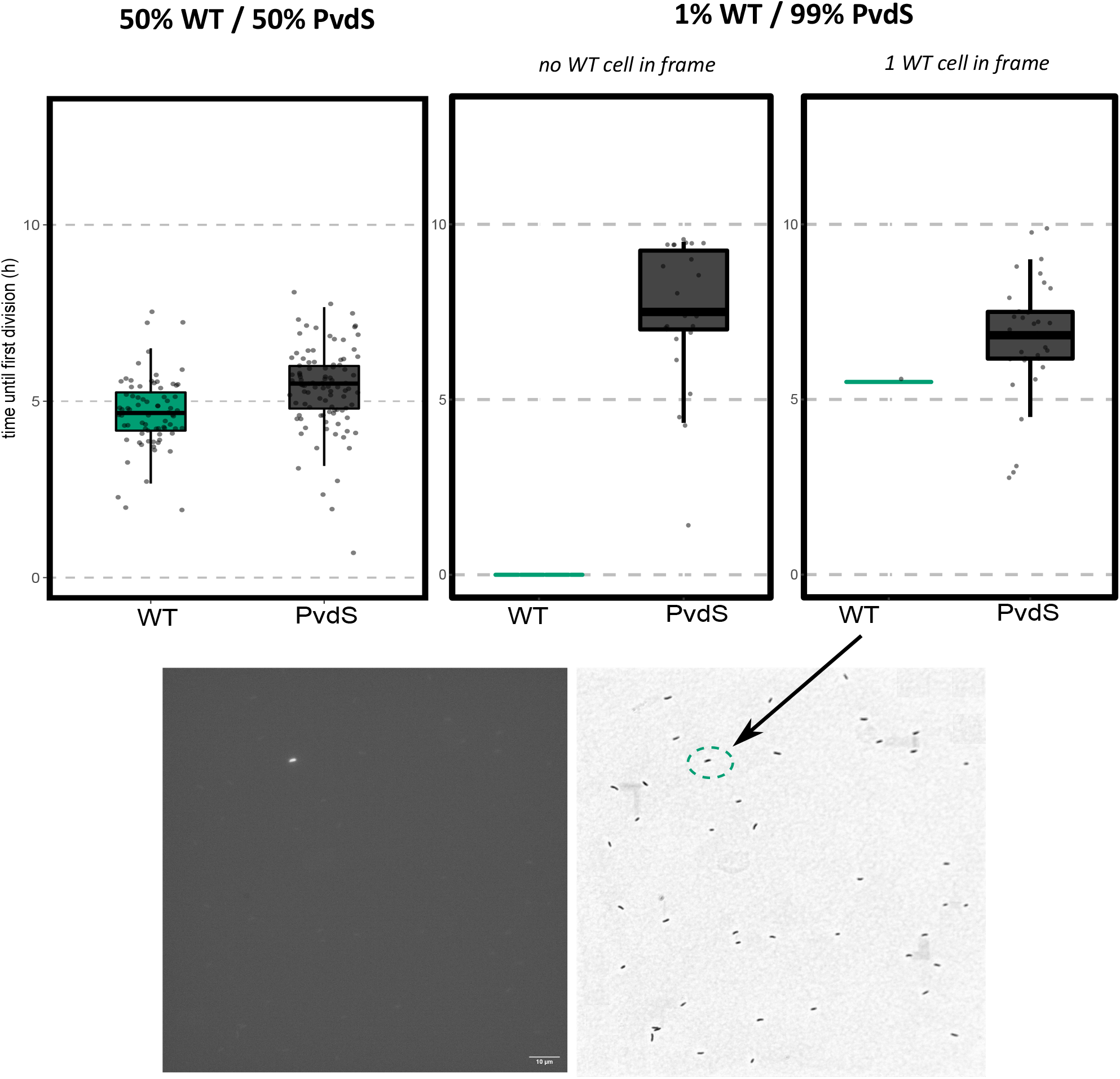
Lag time of ancestor and nonproducing mutant in co-culture where pyoverdin producer is rare. Time after first division of ancestor and mutant cells pre-treated with DP upon co-inoculation on a fresh agarose pad. Prior to inoculation cells were mixed at 1:1 proportions (left panel, figure is the same as fig 4 B in main text and is included for reference) or ancestor cells at 1%. Center panel depicts data from a technical replicate where no ancestor cells can be identified in the microscope field of acquisition, right panel depicts data from a technical replicate where a single ancestor cell can be identified. Images below correspond to this latter dataset, where left panel represents fluorescent pyoverdin imaging and right panel represents phase contrast imaging. Size of the field of view is 132.06 × 132.06 *µ*m and scale bar represents 10*µ*m The single pyoverdin producer is highlighted in green dashed line.(N = 27, 35 respectively)

## References

1. Hans-Curt Flemming, Jost Wingender, Ulrich Szewzyk, Peter Steinberg, Scott A. Rice, and Staffan Kjelleberg. Biofilms: an emergent form of bacterial life. Nature Reviews Microbiology, 14(9):563–575, 9 2016.

2. John W. Newman, Rachel V. Floyd, and Joanne L. Fothergill. The contribution of Pseudomonas aeruginosa virulence factors and host factors in the establishment of urinary tract infections. FEMS Microbiology Letters, 364(15), 8 2017.

3. Médéric Diard, Victor Garcia, Lisa Maier, Mitja N. P. Remus-Emsermann, Roland R. Regoes, Martin Ackermann, and Wolf-Dietrich Hardt. Stabilization of cooperative virulence by the expression of an avirulent phenotype. Nature, 494(7437):353–356, 2 2013.

4. N. R. Gilkes, D. G. Kilburn, R. C. Miller, and R. A.J. Warren. Bacterial cellulases. Bioresource Technology, 36(1):21–35, 1 1991.

5. J. M. Meyer and M. A. Abdallah. The fluorescent pigment of Pseudomonas fluorescens: Biosynthesis, purification and physicochemical properties. Journal of General Microbiology, 107(2), 1978.

6. J. B. Neilands. Iron absorption and transport in microorganisms. Annual review of nutrition, 1, 1981.

7. Michael T Ringel and Thomas Brüser. The biosynthesis of pyoverdines. Microbial cell, 5(10):424–437, 8 2018.

8. Anne Marie Albrecht-Gary, Sylvie Blanc, Natacha Rochel, Aydin Z. Ocaktan, and Mohamed A. Abdallah. Bacterial iron transport: coordination properties of pyoverdin PaA, a peptidic siderophore of Pseudomonas aeruginosa. Inorganic Chemistry, 33(26), 1994.

9. Anne Bonneau, Béatrice Roche, and Isabelle J. Schalk. Iron acquisition in Pseudomonas aeruginosa by the siderophore pyoverdine: an intricate interacting network including periplasmic and membrane proteins. Scientific Reports, 10(1):120, 12 2020.

10. Stuart A. West, Stephen P. Diggle, Angus Buckling, Andy Gardner, and Ashleigh S. Griffin. The social lives of microbes. Annual Review of Ecology, Evolution, and Systematics, 38(1):53–77, 2007.

11. Angus Buckling, Freya Harrison, Michiel Vos, Michael A. Brockhurst, Andy Gardner, Stuart A. West, and Ashleigh Griffin. Siderophore-mediated cooperation and virulence in Pseudomonas aeruginosa. FEMS Microbiology Ecology, 62(2):135–141, 11 2007.

12. Paul B. Rainey, Nicolas Desprat, William W. Driscoll, and Xue-Xian Zhang. Microbes are not bound by sociobiology: Response to Kümmerli and Ross-Gillespie (2013). Evolution, 68(11):3344–3355, 2014.

13. Xue-Xian Zhang and Paul B. Rainey. Exploring the sociobiology of pyoverdin-producing Pseudomonas. Evolution, 67(11):3161–3174, 11 2013.

14. Thomas Julou, Thierry Mora, Laurent Guillon, Vincent Croquette, Isabelle J Schalk, David Bensimon, and Nicolas Desprat. Cell-cell contacts confine public goods diffusion inside Pseudomonas aeruginosa clonal microcolonies. Proceedings of the National Academy of Sciences of the United States of America, 110(31):12577–82, 7 2013.

15. Zhenyu Jin, Jiahong Li, Lei Ni, Rongrong Zhang, Aiguo Xia, and Fan Jin. Conditional privatization of a public siderophore enables Pseudomonas aeruginosa to resist cheater invasion. Nature Communications, 9(1):1383, 12 2018.

16. Susse Kirkelund Hansen, Paul B. Rainey, Janus A.J. Haagensen, and Søren Molin. Evolution of species interactions in a biofilm community. Nature, 445(7127), 2007.

17. Otto X. Cordero and Manoshi S. Datta. Microbial interactions and community assembly at microscales. Current Opinion in Microbiology, (31):227–234, 2016.

18. Alma Dal Co, Simon van Vliet, Daniel Johannes Kiviet, Susan Schlegel, and Martin Ackermann. Short-range interactions govern the dynamics and functions of microbial communities. Nature Ecology and Evolution, 4(3):366–375, 3 2020.

19. Raimo Hartmann, Praveen K. Singh, Philip Pearce, Rachel Mok, Boya Song, Francisco Díaz-Pascual, Jörn Dunkel, and Knut Drescher. Emergence of three-dimensional order and structure in growing biofilms. Nature Physics, 15(3):251–256, 3 2019.

20. Mark W. Silby, Ana M. Cerdeño-Tárraga, Georgios S. Vernikos, Stephen R. Giddens, Robert W. Jackson, Gail M. Preston, Xue Xian Zhang, Christina D. Moon, Stefanie M. Gehrig, Scott A.C. Godfrey, Christopher G. Knight, Jacob G. Malone, Zena Robinson, Andrew J. Spiers, Simon Harris, Gregory L. Challis, Alice M. Yaxley, David Harris, Kathy Seeger, Lee Murphy, Simon Rutter, Rob Squares, Michael A. Quail, Elizabeth Saunders, Konstantinos Mavromatis, Thomas S. Brettin, Stephen D. Bentley, Joanne Hothersall, Elton Stephens, Christopher M. Thomas, Julian Parkhill, Stuart B. Levy, Paul B. Rainey, and Nicholas R. Thomson. Genomic and genetic analyses of diversity and plant interactions of Pseudomonas fluorescens. Genome Biology, 10(5):R51, 5 2009.

21. Christina D Moon, Xue-Xian Zhang, Sandra Matthijs, Mathias Schäfer, Herbert Budzikiewicz, and Paul B Rainey. Genomic, genetic and structural analysis of pyoverdine-mediated iron acquisition in the plant growthpromoting bacterium Pseudomonas fluorescens SBW25. BMC Microbiology, 8(1):7, 2008.

22. Katrin Hammerschmidt, Caroline J Rose, Benjamin Kerr, and Paul B Rainey. Life cycles, fitness decoupling and the evolution of multicellularity. Nature, 515(7525):75–9, 11 2014.

23. A. J. Spiers, S. G. Kahn, J. Bohannon, M. Travisano, and P. B. Rainey. Adaptive divergence in experimental populations of Pseudomonas fluorescens. I. Genetic and phenotypic bases of wrinkly spreader fitness. Genetics, (161):33–46, 2002.

24. Handuo Shi, Corey S. Westfall, Jesse Kao, Pascal D. Odermatt, Sarah E. Anderson, Spencer Cesar, Montana Sievert, Jeremy Moore, Carlos G. Gonzalez, Lichao Zhang, Joshua E. Elias, Fred Chang, Kerwyn Casey Huang, and Petra Anne Levin. Starvation induces shrinkage of the bacterial cytoplasm. Proceedings of the National Academy of Sciences, 118(24):e2104686118. 6 2021.

25. Emilie Yeterian, Lois W. Martin, Laurent Guillon, Laure Journet, Iain L. Lamont, and Isabelle J. Schalk. Synthesis of the siderophore pyoverdine in Pseudomonas aeruginosa involves a periplasmic maturation. Amino Acids, 38(5):1447–1459, 5 2010.

26. Emilie Yeterian, Lois W. Martin, Iain L. Lamont, and Isabelle J. Schalk. An efflux pump is required for siderophore recycling by Pseudomonas aeruginosa. Environmental Microbiology Reports, 2(3):412–418, 12 2009.

27. Isabelle J. Schalk, Christophe Hennard, Christophe Dugave, Keith Poole, Mohamed A. Abdallah, and Franc Pattus. Iron-free pyoverdin binds to its outer membrane receptor FpvA in Pseudomonas aeruginosa: a new mechanism for membrane iron transport. Molecular Microbiology, 39(2):351–361, 1 2001.

28. Isabelle J. Schalk and Laurent Guillon. Pyoverdine biosynthesis and secretion in Pseudomonas aeruginosa : implications for metal homeostasis. Environmental Microbiology, 15(6):1661–1673, 6 2013.

29. Maxime Ardré, Djinthana Dufour, and Paul B. Rainey. Causes and biophysical consequences of cellulose production by pseudomonas fluorescens sbw25 at the air-liquid interface. Journal of Bacteriology, 201(18), 9 2019.

30. Michael J McDonald, Stefanie M Gehrig, Peter L Meintjes, Xue-Xian Zhang, and Paul B Rainey. Adaptive divergence in experimental populations of Pseudomonas fluorescens. IV. Genetic constraints guide evolutionary trajectories in a parallel adaptive radiation. Genetics, 183(3):1041–53, 11 2009.

31. P.R.J. Yulo, N. Desprat, M.L. Gerth, Y. Liu, X.X. Zhang, P.B. Rainey, and H.L. Hendrickson. Experimental Evolution of Cell Shape in Bacteria. bioRxiv, page 263681, 2 2018.

32. Steven D. Quistad, Guilhem Doulcier, and Paul B. Rainey. Experimental manipulation of selfish genetic elements links genes to microbial community function. Philosophical Transactions of the Royal Society B: Biological Sciences, 375(1798):20190681, 5 2020.

33. Xue-Xian Zhang, Stephen R Ritchie, Hao Chang, Dawn L Arnold, Robert W Jackson — Paul, and B Rainey. Genotypic and phenotypic analyses reveal distinct population structures and ecotypes for sugar beet-associated Pseudomonas in Oxford and Auckland. Ecology and Evolution, 10:5963–5975, 2020.

34. Pierre Cornelis and Sandra Matthijs. Diversity of siderophore-mediated iron uptake systems in fluorescent pseudomonads: Not only pyoverdines. Environmental Microbiology, 4(12):787–798, 12 2002.

35. Jeff Gore, Hyun Youk, and Alexander van Oudenaarden. Snowdrift game dynamics and facultative cheating in yeast. Nature, 459(7244):253–6, 5 2009.

36. Rebecca L. Scholz and E. Peter Greenberg. Sociality in Escherichia coli: Enterochelin is a private good at low cell density and can be shared at high cell density. Journal of Bacteriology, 197(13):2122–2128, 2015.

37. Lucy Shapiro, Harley H. McAdams, and Richard Losick. Generating and exploiting polarity in bacteria. Science, 298(5600):1942–1946, 12 2002.

38. David M. Raskin and Piet A.J. De Boer. Rapid pole-to-pole oscillation of a protein required for directing division to the middle of Escherichia coli. Proceedings of the National Academy of Sciences of the United States of America, 96(9):4971, 4 1999.

39. Géraldine Laloux and Christine Jacobs-Wagner. How do bacteria localize proteins to the cell pole? Journal of Cell Science, 127(1), 2013.

40. Laurent Guillon, Maher El Mecherki, Stephan Altenburger, Peter L. Graumann, and Isabelle J. Schalk. High cellular organization of pyoverdine biosynthesis in Pseudomonas aeruginosa: clustering of PvdA at the old cell pole. Environmental Microbiology, 14(8):1982–1994, 8 2012.

41. Véronique Gasser, Laurent Guillon, Olivier Cunrath, and Isabelle.J. Schalk. Cellular organization of siderophore biosynthesis in Pseudomonas aeruginosa: Evidence for siderosomes. Journal of Inorganic Biochemistry, 148:27–34, 2015.

42. Özhan Özkaya, Roberto Balbontín, Isabel Gordo, and Karina B Xavier. Cheating on cheaters stabilizes cooperation in Pseudomonas aeruginosa. Current Biology, 28(13):2070–2080, 7 2018.

43. Philippe Remigi, Gayle C. Ferguson, Ellen McConnell, Silvia De Monte, David W. Rogers, and Paul B. Rainey. Ribosome provisioning activates a bistable switch coupled to fast exit from stationary phase. Molecular Biology and Evolution, 36(5):1056–1070, 5 2019.

44. Joseph W Kloepper, John Leongt, Martin Teintzet, and Milton N Schroth. Enhanced plant growth by siderophores produced by plant growth-promoting rhizobacteria. Nature, 286:885–886, 1980.

45. Dieter Haas and Geneviève Défago. Biological control of soil-borne pathogens by fluorescent pseudomonads. Nature Reviews Microbiology, 3(4):307–319, 4 2005.

46. Peter Stilwell, Chris Lowe, and Angus Buckling. The effect of cheats on siderophore diversity in Pseudomonas aeruginosa. Journal of Evolutionary Biology, 31(9):1330– 1339, 9 2018.

47. Pascal Mirleau, Sandrine Delorme, Laurent Philippot, Jean-Marie Meyer, Sylvie Mazurier, and Philippe Lemanceau. Fitness in soil and rhizosphere of Pseudomonas fluorescens C7R12 compared with a C7R12 mutant affected in pyoverdine synthesis and uptake. FEMS Microbiology Ecology, 34(1):35–44, 10 2000.

48. Ehud Banin, Michael L Vasil, and E Peter Greenberg. Iron and Pseudomonas aeruginosa biofilm formation. Proceedings of the National Academy of Sciences, 102(31):11076–11081, 2005.

49. Gabriele Sass, Hasan Nazik, John Penner, Hemi Shah, Shajia Rahman Ansari, Karl V. Clemons, Marie Christine Groleau, Anna Maria Dietl, Paolo Visca, Hubertus Haas, Eric Déziel, and David A. Stevens. Studies of Pseudomonas aeruginosa mutants indicate pyoverdine as the central factor in inhibition of Aspergillus fumigatus biofilm. Journal of Bacteriology, 200(1), 1 2018.

50. Adrienne M. Brauer, Handuo Shi, Petra Anne Levin, and Kerwyn Casey Huang. Physiological and regulatory convergence between osmotic and nutrient stress responses in microbes. Current Opinion in Cell Biology, 81:102170, 4 2023.

51. Xue-Xian Zhang and Paul B Rainey. Genetic analysis of the histidine utilization (hut) genes in Pseudomonas fluorescens SBW25. Genetics, 176(4):2165–76, 8 2007.

52. Xue-Xian Zhang and Paul B. Rainey. Construction and validation of a neutrally-marked strain of Pseudomonas fluorescens SBW25. Journal of Microbiological Methods, 71(1):78–81, 10 2007.

53. Stella Stylianidou, Connor Brennan, Silas B. Nissen, Nathan J. Kuwada, and Paul A. Wiggins. SuperSegger: robust image segmentation, analysis and lineage tracking of bacterial cells. Molecular Microbiology, 102(4):690–700, nov 2016.

54. W.S Rasband. ImageJ. U. S. National Institutes of Health, Bethesda, https://imagej.nih.gov/ij/. Maryland, USA, 1997-2021.

55. MATLAB. version 9.5.0.1265761 (R2018b) Update 6). The MathWorks Inc., Natick, Massachusetts, 2018.

56. MATLAB. Mastering Machine Learning -A Step-by-Step Guide with MATLAB. MathWorks, 2019.

57. R Core Team. R: A Language and Environment for Statistical Computing. R Foundation for Statistical Computing, Vienna, Austria, 2020. https://www.R-project.org/.

58. Kyoung Hee Choi, Jared B. Gaynor, Kimberly G. White, Carolina Lopez, Catharine M. Bosio, Rox Ann R. Karkhoff-Schweizer, and Herbert P. Schweizer. A Tn7-based broadrange bacterial cloning and expression system. Nature methods, 2(6):443–448, jun 2005.

59. Rudolf O. Schlechter, Hyunwoo Jun, Micha-l Bernach, Simisola Oso, Erica Boyd, Dian A. Muñoz-Lintz, Renwick C.J. Dobson, Daniela M. Remus, and Mitja N.P. Remus-Emsermann. Chromatic bacteria – A broad host-range plasmid and chromosomal insertion toolbox for fluorescent protein expression in bacteria. Frontiers in Microbiology, 9(DEC):423623, dec 2018.

60. Chaokun Li, Aiyun Wen, Benchang Shen, Jia Lu, Yao Huang, and Yongchang Chang. FastCloning: A highly simplified, purification-free, sequence- and ligation-independent PCR cloning method. BMC Biotechnology, 11(1):1–10, oct 2011.

61. Tsuyoshi Uehara, Thuy Dinh, and Thomas G. Bernhardt. LytM-domain factors are required for daughter cell separation and rapid ampicillin-induced lysis in Escherichia coli. Journal of bacteriology, 191(16):5094–5107, aug 2009.

62. Ying Bao, Douglas P. Lies, Haian Fu, and Gary P. Roberts. An improved Tn7-based system for the single-copy insertion of cloned genes into chromosomes of gram-negative bacteria. Gene, 109(1):167–168, ec 1991.

